# Antiviral and Antibacterial Efficacy of Nanocomposite Amorphous Carbon Films with Copper Nanoparticles

**DOI:** 10.1101/2023.10.13.562202

**Authors:** Shahd Bakhet, Asta Tamulevičienė, Andrius Vasiliauskas, Mindaugas Andrulevičius, Šarūnas Meškinis, Sigitas Tamulevičius, Neringa Kašėtienė, Mindaugas Malakauskas, Raimundas Lelešius, Dainius Zienius, Algirdas Šalomskas, Krišjānis Šmits, Tomas Tamulevičius

## Abstract

Copper compound-rich films and coatings are effective against widespread viruses and bacteria. Even though the killing mechanisms are still debated it is agreed that the metal ion, nanoparticle release, and surface effects are of paramount importance in the antiviral and antibacterial efficacy of the surfaces. In this work we have investigated the behaviour of the reactive magnetron sputtered nanocomposite diamond-like carbon thin films with copper nanoparticles (DLC:Cu). The films were etched employing oxygen plasma and/or exposed to ultra-pure water aiming to investigate the differences of the Cu release in the medium and changes in film morphology. The presence of metallic copper and Cu_2_O phases were confirmed by multiple analytical methods. Pristine films were more effective in the Cu release reaching up to 1.3 mg/L/cm^2^ concentration. Plasma processing resulted in the oxidation of the films which released less Cu but after exposure to water, their average roughness increased more, up to 5.5 nm. Pristine and O_2_ plasma processed DLC:Cu films were effective against both model coronavirus and herpesvirus after 1-hour contact time and reached virus reduction up to 2.23 and 1.63 log_10_, respectively. Pristine DLC:Cu films were more effective than plasma-processed ones against herpesvirus, while less expressed difference was found for coronavirus. The virucidal efficacy over up to 24 h exposures in the aqueous medium was validated. A bactericidal study confirmed that pristine DLC:Cu films were effective against gram-negative *E. coli* and gram-positive *E. faecalis* bacteria. After 3 hours 100% antibacterial efficiency (ABE) was obtained for *E. coli* and 99.97% for *E. faecalis*. After 8 hours and longer exposures, 100% ABE was reached. The half-life inactivation of viruses was 8.10 – 11.08 minutes and for *E. faecalis* 15.1 – 72.2 minutes.

## 1. INTRODUCTION

In recent years, the emergence and spread of antibiotic-resistant bacteria have posed significant challenges to healthcare and public health systems worldwide ^1^. Moreover, the recent COVID-19 pandemic illustrated the importance and need for efficient antiviral surfaces, especially on high-touch surfaces ^2–4^. Intense research efforts have been applied to develop efficient antibacterial strategies. Among the various approaches explored, the utilization of novel materials with inherent antibacterial properties has shown great promise. It has been shown that silver, copper, and their compounds are among the most popular materials used for different antimicrobial and biomedicine applications ^5, 6^ because of their high bactericidal performance ^7^. It should be noted that Cu has been recognized by the United States Environmental Protection Agency (EPA) as the first metallic antimicrobial agent in 2008 but with ongoing waterborne hospital-acquired infections and antibiotic resistance, research on Cu as an antimicrobial agent is booming again ^8^. It is commonly proven that the antimicrobial properties of metals depend very much on their size and shape, and they are most efficient when the metal is in the form of nanoparticles (NPs). Recent studies on Ag and Cu NPs have shown that there is a strong influence of metal particle size, shape, surface charge, and other factors on the release of metal ions that are critical to bactericidal and virucidal properties. Smaller metal NPs oxidize easier, releasing more metal ions which in turn increases the bactericidal effect ^7^. Cu is less expensive than Ag, but it is also a very practical choice because it possesses not only bactericidal but also virucidal properties at the same time and can serve as a multifunctional protecting coating ^9, 10^. Applications of metal ions and NPs require their entrapment of other materials such as polymers, ceramics, metals, and composites, where they are held in place by different means and thereby augment the material with antibacterial activity^11^.

Different approaches are used to produce nanocomposite antimicrobial surfaces with embedded metal nanoparticles where vacuum deposition methods provide excellent controllability and repeatability of the process ^12^. One such material widely used by researchers is amorphous diamond-like (DLC) nanocomposite coating with IB group metal nanoparticles (Au, Ag, Cu), which combines the desirable mechanical properties of DLC and the bactericidal properties of metal nanoparticles. DLC is an amorphous carbon (a-C) material that exhibits exceptional mechanical, chemical, and biocompatible properties, making it an attractive candidate for various applications. ^13^. On the other hand, metal nanoparticles, and especially Cu have been extensively studied for their potential for antimicrobial activity. By combining these two materials, the amorphous diamond-like carbon-copper (DLC:Cu) nanocomposite coatings offer a unique opportunity to enhance the antibacterial properties beyond what each material can achieve individually ^5^.

Probably the most popular way to produce such nanocomposites is different modes of magnetron (either reactive magnetron) sputtering where different targets (or different hydrocarbon gases) are applied. This versatile method enables the production of coatings with a controllable composition resulting in efficient bactericidal and other (optical, electrical, mechanical) properties ^14–18^. An alternative for Cu NPs introduction in the a-C films is cluster beam deposition combined with plasma-enhanced chemical vapour deposition^19^.

The metal filler loading in the amorphous carbon film has an impact not only on its biocompatibility or antimicrobial properties but also on the internal bonding as it was confirmed by Raman spectroscopy in different studies indicating increasing graphitic *sp*^2^ boding over *sp*^3^ diamond phase. That relates to the optical, electrical, tribological, hardness, biofouling, and adhesion properties of the DLC films^20–22^. Moreover, metal loading reduces the residual stress and the mismatch of interfacial atomic bonding increasing the adhesion of the films on different metallic and non-metallic substrates^23–25^.

The efficacy of the metal nanoparticle films and coatings depends on their ability to release the metal ions and nanoparticles in the aqueous media. It was reported that DLC:Cu ^26^, plasma-processed DLC:Ag nanocomposite films ^27^, and ultra nanocrystalline diamond protective films on Ag nanoparticles formed by the dewetting process release metal ions in aqueous media after days of immersion in water ^12^. Studies of polyethylene nanocomposites with Ag or Cu nanoparticles indicated significant differences in the absolute amount of metal release Cu being hundreds of times more effective ^28^. Metal release studies were not limited to ultra-pure water but included immersion in standard simulated body fluid ^29^, artificial seawater ^26^, and different pH biological solutions ^30, 31^. The release of Cu NPs and Cu ions depends on the coating structure^19^, contact time, and amount of the solvent (medium)^32, 33^ where it can lead to oxidation and fast degradation of metallic Cu resulting in the burst release happening at the first immersion stages. This may affect the antibacterial and antiviral properties and have cytotoxic effect^19^. Therefore, researching the coating compositions demonstrating slow and controlled Cu release is of paramount importance ^34^. In most cases, the metal release follows saturation behaviour over time and the release is understood not only as a surface phenomenon but also governed by a diffusion process in which the active metal entities must move from the interior of the coating specimen to the surface to be released ^35^.

In our previous work, we have demonstrated that DLC:Ag films are effective against *Campylobacter jejuni* and *Listeria monocytogenes* on food preparation surfaces ^36^ and *Staphylococcus aureus* including methicillin-resistant strains aiming for development of antimicrobial bandages ^27^ and patches ^37^. While gentle oxygen plasma processing was essential for the high antibacterial efficacy of DLC:Ag films ^27, 37^ use of a harsh CF_4_/O_2_ mixture resulted in the removal of DLC matrix leaving only the agglomerated Ag filler ^38^.

Despite the broad range of DLC:Cu antibacterial studies there is still a very limited number of works investigating antiviral applications of such coatings ^19, 39^. Moreover, the combined antiviral and antimicrobial behaviour is often obtained using more complex compositions like, for example, Ag and Cu alloys ^10^ while in this work Cu metal filler was confirmed efficient against both types of germs alone.

In this work, we demonstrate that reactive high-power impulse magnetron sputtered nanocomposite amorphous diamond-like carbon thin films with copper nanoparticles have expressed not only the virucidal properties against animal-derived model viruses, including ribonucleic acid (RNA) containing coronavirus and deoxyribonucleic acid (DNA) containing herpesvirus but at the same time they demonstrate antibacterial effect against gram-negative *Escherichia coli* and gram-positive *Enterococcus faecalis* bacteria. The detailed studies of the Cu release in aqueous media, surface morphology evolution, and elemental changes of oxygen plasma processed, and ultra-pure water-exposed films helped in understanding their possible antibacterial and antiviral properties.

## 2. MATERIALS AND METHODS

### 2.1. Deposition of DLC Films

#### 2.1.1 Deposition of DLC:Cu Films

The hydrogenated amorphous diamond-like carbon films with copper nanoparticles (DLC:Cu or otherwise a-C:H:Cu) films were deposited on crystalline silicon substrates (p-type (100), MicroChemicals GmbH, Germany) by unbalanced reactive magnetron sputtering of the 3″ diameter pure Cu target (99.99% purity, Testbourne B.V., The Netherlands) using high-power impulse magnetron sputtering (HIPIMS) mode in a custom made vacuum deposition system ^13^. Argon gas was used for sputtering (99.999%, Linde Gass, Lithuania), and acetylene (99.6%, Linde Gass, Lithuania) was used as a reactive gas with 63 sccm and 10 sccm flow rates, respectively. The base pressure was 8·10^-4^ Pa and the work pressure of 7 ± 1·10^-1^ Pa was maintained in the chamber throughout the deposition process. The average magnetron target current was *I*_av_=0.12 A, voltage *V*=687 V, while the pulse current was *I*_pulse_=26 A with on-time *t*_on_=100 µs and off-time *t*_off_=10 000 μs. The substrates were grounded and the distance between magnetron target and the samples was 0.1 m.

The thickness of the nanocomposite films was monitored with a quartz microbalance and resulted in 100 nm.

#### 2.1.2. Deposition of Cu-free DLC Films

Copper-free DLC films for reference virucidal testing were deposited employing a closed drift ion beam source with an annular discharge channel from acetylene gas (99.6%. Linde Gas, Lithuania). The supplied gas is ionized close to the anode due to the discharge channel, and further ions are accelerated in a magnetic field and form a layer on the substrates mounted on the water-cooled cathode. The deposition was performed at 0.7 keV ion beam energy and working pressure 1·10^-2^ Pa. Thickness of the film was set to 170 nm. Deposition resulted in hydrogenated amorphous diamond-like carbon (DLC) films. More details about the deposition procedure can be found in ^40^.

### 2.2. O_2_ plasma processing of the DLC:Cu films

Some of the DLC:Cu samples were dry-etched using a Plasma-600-T (AO Kvarts, USSR) device. The samples were placed inside the processing chamber and were etched by radio frequency (RF; 13.56 MHz) oxygen plasma (99.999%, Linde Gass, Lithuania) in 133 Pa pressure and 0.3 W/cm² power for 5 s or 20 s. The RF oxygen plasma etching was intended for the removal of the thin amorphous DLC carbon layer from the Cu NP surface.

### 2.3. Cu release in aqueous media

Cu release from the pristine and O_2_ plasma processed DLC:Cu films on 0.9x0.9 cm^2^ silicon substrates into ultra-pure water (ChromAR HPLC, pH = 5.0 – 8.0, Macron Fine Chemicals, Avantor, USA) was investigated by immersing samples in 20 ml for different periods of 24 hours and 168 hours and kept at constant temperature of 5°C and 37°C. A preliminary study of Cu release in cell growth media was conducted at room temperature immersing samples for different durations from 5 to 120 minutes, more details are provided in the supplements. The released Cu compound content in the liquids was determined with a double-beam atomic absorption spectrometer (AAS) AAnalyst 400 (Perkin Elmer, USA). The Cu amount was determined from the reference solution calibration curve. The measured optical densities were normalized to the calibration curve obtaining the released Cu concentrations. The concentrations were normalized to the sample surface area.

### 2.4. Physical Characterization of the DLC:Cu Films

Field emission gun scanning electron microscope (SEM) Quanta 200 FEG (FEI, USA) with energy dispersive X-ray spectrometer (EDS) XFlash 4030 (Bruker, Germany) was used for the imaging and surface composition analysis of DLC:Cu samples, respectively, after the deposition as well as RF oxygen plasma etching and immersion experiments. 10 kV accelerating voltage was used for the microanalysis and the obtained results were averaged from three measurements.

Focused ion beam (FIB) milling was used in SEM Helios NanoLab 650 (FEI, Netherlands) and Helios 5 UX (Thermo Scientific, MA, USA) for preparation of the DLC:Cu thin film sample lamella for cross-section inspection under transmission electron microscope (TEM) Tecnai G2 F20 X-TWIN (FEI, Netherlands) with a Schottky type field emission electron source operated at 200 kV. Selected area electron diffraction (SAED) was analysed using CrysTBox software^42^.

The atomic force microscope (AFM) NanoWizard®^3^ NanoScience (JPK instruments AG, Germany) operating in a tapping mode was used for the quantitative studies of the roughness of the pristine and water-immersed samples. An I-shaped silicon cantilever with a 13–77 N/m spring constant and a 10.0 nm typical tip curvature radius was used for the measurements.

The structure and carbon phases of the DLC matrix were probed employing a micro Raman scattering spectrometer inVia (Renishaw, United Kingdom) that is equipped with a 532 nm solid-state laser, x50 objective. Each sample was measured twice at two different locations, the final Raman spectrum of a sample is the mean result of 3 scans. The laser power on the sample was 1.75 mW, integration time was set to 10 s. The Raman Stokes signal was dispersed with a diffraction grating (2400 grooves/mm) and data was registered using a Peltier-cooled charge-coupled device (CCD) 1024×256 pixel detector. Silicon was used to calibrate the Raman setup in both Raman wavenumber and spectral intensity. Analysis of spectra was performed to reveal *sp*^2^ and *sp*^3^ carbon phases in the DLC matrix of the nanocomposite films. Peaks were fitted using two Gaussian functions and characteristic parameters such as D and G peak intensity ratio as well as G peak position and full width at half maximum (FWHM) were revealed.

The chemical composition and crystal structures of the pristine DLC:Cu films were carried out using a D8 Advance X-ray diffractometer (Bruker, Germany) operating at 40 kV and 40 mA with Cu K_α_ (λ = 1.5418 Å) at grazing angle arrangement. More details can be found in^41^.

Chemical composition and chemical bonds of the elements in the pristine and differently exposed DLC:Cu films was studied by X-Ray Photoelectron Spectroscopy method employing a XSAM800 (Kratos Analytical, UK) spectrometer. The hemispherical electron energy analyser was in the fixed transition mode (FAT), pass energy was set to 20 eV and energy increment was set to 0.1 eV. Non-monochromatic Al Kα radiation (*hν* = 1486.6 eV) was used as excitation source..

### 2.5 Abrasion Test

The DLC:Cu samples were tested with the custom-made abrasion device similar to that described by Mehanna *et al.* ^43^. The samples were placed on the rotating holder that was in physical contact with the secured 2 mm diameter sandpaper of 2500, 1000 and 400 grit. Samples were rotated with 50 and 100-revolution speeds per minute and experienced 10, 20, or 30 rotation cycles. Pristine surface and after mechanical impact the samples were inspected under an optical microscope B-600MET (Optika, Italy) with a 3.2 MPix CMOS digital camera Optical Pro 3 (Optika, Italy).

### 2.6. Antiviral Testing

The antiviral testing covered pristine DLC:Cu films, 5 s processed DLC:Cu films, and Cu free DLC films as well as Si substrate for control.

#### 2.6.1. Model Viral Strains Used and Cultivation of Cells

The avian infectious bronchitis virus (IBV) Beaudette strain was obtained from Dr. M. H. Verheije (Utrecht University, The Netherlands). The agent of bovine herpesvirus infection virus (BoHV-1) 4016 strain was provided by Dr. I. Jacevičienė (National Food and Veterinary Risk Assessment Institute, Lithuania).

Vero (ATCC CCL-81™) and MDBK (NBL-1, Bov.90050801) cell lines were obtained from Dr. I. Jacevičienė. Vero and MDBK/NBL-1 cells were cultured in Dulbecco’s modified eagle’s medium (DMEM, Gibco, UK) and DMEM/Nutrient Mixture F-12 (DMEM / F-12, Gibco, UK), respectively with 10% fetal bovine serum (FBS, Gibco, UK) at 37°C in a 5% CO_2_ incubator. Nystatin (100 units/ml) and Gentamicin (50 μg/ml) were used to prevent microbial contamination for Vero cells, while Penicillin (100 units/ml) and Streptomycin (100 μg/ml) were used to prevent microbial contamination for MDBK/NBL-1 cell. The tests were performed at temperatures of 20±2°C and 37±1°C depending on the method used.

#### 2.6.2. Cytotoxicity Control

The MDBK and Vero cells were cultured in a 96-well plate and the medium culture was replaced with 0.1 ml of tested DLC:Cu film extract as a negative control, Triton X-100 (Sigma Aldrich, MA, USA) as a positive control and blank, respectively. The test and control (negative and positive) samples were prepared according to the recommendations^44^. The most sensitive extract test was chosen for the cytotoxicity assessment of the coatings^45^. Cultures were then incubated for 24 ± 1 h at 37 ± 1°C in a humidified atmosphere containing 5% CO_2_ and 95% air. After 24 ± 1 h of incubation, cell morphology in each well was assessed microscopically and scored from 0 to 4 grade as described in ISO 10993-5 standard^46^. Cells were inspected employing an inverted optical microscope DMiL (Leica) equipped with a 3.1 Mpixel camera Optikam Pro 3 (Optika). Clarity and pH of the cell culture medium were also assessed visually^47^ and a colorimetric cell viability MTT assay^48^ was performed.

Cytotoxic concentration (CC_50_) was calculated by using Quest Graph™ IC_50_ Calculator and represents the concentration at which a substance exerts half of its maximal inhibitory effect.

#### 2.6.3. Virucidal Efficacy Testing

##### 2.6.3.1 Virus Treatment

The potential effectiveness of the virucidal effect of the DLC:Cu and DLC thin films and the change in viral load was evaluated by determining the tissue culture infectious dose (TCID_50_) of intact viruses (control) and after their exposure (affected viruses). Suspensions of coronaviruses and herpesviruses in 0.01 ml were incubated for 1 h at room temperature with investigated samples. Cultivation and propagation of IBV was performed by infecting Vero cells while BoHV-1 strain 4016 was cultivated in MDBK/NBL-1 cell culture ^49^.

##### 2.6.3.2 Virus Quantification

Virucidal efficiency was evaluated by comparing changes in the concentration of viruses before and after contact with the surface of DLC:Cu and DLC coatings. After 48-120 hours of incubation, the cytopathogenic effect (CPE) in cells was assessed with an optical microscope. Viral titre, mean and standard deviation (*M*±*σ*), were estimated by the Spearman-Karber method ^50, 51^. The half-life of the viruses was calculated employing **eq. S1**.

##### 2.6.3.3 Viral Nucleic Acid Quantification

For IBV real-time TaqMan reverse transcription polymerase chain reaction (PCR) was performed according to Meir ^52^. The sequences of the conserved region of the N gene of strain IBV H120 at nucleotide positions 741–1077, consisting of 336 base pairs, were used to design primers (GenBank no. AM260960). For the reaction, StepOne™ Plus real-time PCR mix (Applied Biosystems, UK) was used. Amplification plots were evaluated, and threshold cycle (Ct) values were determined using a Mastercycler RealPlex2 (Eppendorf, Germany). The reaction is carried out at 45°C for 10 minutes and 95°C for 10 minutes, 40 cycles consisting of a denaturation step at 95°C for 15 s, and an elongation step at 60°C for 45 s.

For BoHV-1 real-time PCR was performed according to Abril *et al.* ^53^. BoHV-1 glycoprotein B gene sequences were amplified for quantitative real-time PCR (TaqMan). Oligonucleotide primers and MGB-labelled probes were synthesized by Invitrogen (USA). Amplifications were performed using TaqMan Universal Master Mix II (catalogue no. 4440038, Applied Biosystems, USA). Mastercycler RealPlex2 (Eppendorf, Germany) was used to record amplification plots that analysed for determining Ct. Isolation of DNA was performed using the Genomic DNA purification kit (K0721, Thermo Fisher Scientific, USA) following the manufacturer’s instructions. Primers and probes for quantitative real-time PCR were generated with Primer Express equipment (version 1.0, Applied Biosystems, Foster City, CA) to amplify BoHV-1 glycoprotein B sequences with an open reading frame (ORF). Oligonucleotide primers and MGB-labelled probes were synthesized by Invitrogen (USA). Extensions were performed according to a standard protocol. PCR reaction conditions: 2 min at 50°C, 10 min at 95°C, followed by 40 cycles consisting of a denaturation step at 95°C for 15 s and an elongation step at 60°C for 1 min.

##### 2.6.3.4 Virucidal Efficacy Stability Test

For determination of DLC:Cu film virucidal efficacy stability, or in short, durability, samples were in continuous contact with virus or medium. Four cycles of treatment with virus and three cycles of treatment with medium were carried out at room temperature. The surfaces were treated with viruses at the beginning of experiment, after 1, 3, 8, and 24 h of incubation, every cycle of virus treatment was followed by the cycle of treatment with medium. The duration of every cycle of virus treatment was 1 hour, while the duration of the first, the second and the third cycles of treatment with medium took 2, 4 and 15 hours, respectively. The experiment is visualized in the supplementary **Table S1.**

The experiments were repeated twice. For the treatment with virus, the suspensions of coronaviruses (5.00±0.19 log_10_ TCID_50_) and herpesviruses (7.25±0.25 log_10_ TCID_50_) in 0.01 ml were placed on the surface of coatings in 24-well plates. After incubation the samples were submerged in 490 μL/well with medium and used for determination of virus titer. For the treatment of medium these samples of coatings were rinsed with 500 μL of Phosphate-Buffered Saline (PBS) and submerged again in 500 μL of medium.

### 2.7. Antibacterial Testing

The antimicrobial efficacy of the DLC:Cu, DLC films and Si substrates was tested employing ATCC 25922 strain of gram-negative *Escherichia coli* and ATCC 29212 strain of gram-positive *Enterococcus faecalis* bacteria.

The bacteria were kept in -80°C brain-heart infusion (BHI, Oxoid, UK) broth with a 30% glycerol supplement before the test. *E. coli* and *E. faecalis* bacteria were inoculated on Lysogeny (LB, Liofilchem, Italy) agar and grown for 24 h at 35°C. PBS broth was used to prepare the suspension (10^6^ colony forming units (CFU)/ml). The pristine DLC:Cu films were disinfected in 70% w/w ethanol before the test, dried in a biosafety cabinet, and then sterilized with a UV lamp for 24 hours. Sterile samples were placed in a 24-well plate, 1 ml of *E. coli* or *E. faecalis* bacterial suspension was poured on them and 1 ml *of E. coli* and *E. faecalis* bacterial culture suspensions were incubated in wells without the samples as control. The samples were incubated for 24 h at 35°C. After 1, 3, 8, and 24 hours, a 100 µl sample was taken from each well. Tenfold dilutions (up to 10^-3^ in PBS broth) were prepared and plated on LB agar plates. The inoculations were repeated three times. The plates were incubated for 24 hours at 35°C. The antibacterial activity (ABE, %) of DLC:Cu films against *E. coli* and *E. faecalis* bacteria was calculated normalizing to the reference sample (**eq. S2**) after 1, 3, 8, and 24 hours. The half-life employing **eq. S1**. The tests were repeated three times.

### 2.8. Statistical Analysis

The experimental results were analysed with OriginPro 2015 (OriginLab) and the mean comparative studies of TCID_50_, and Ct estimation were done by two-way analysis of variance (ANOVA) with the quantitative data expressed as the *M*±*σ*. A statistical significance was considered at \**p* <0.05.

## 3. RESULTS

### 3.1 Elemental and Structural Properties of Pristine DLC:Cu Films

Reactive magnetron sputtered DLC:Cu films had nanoparticles visible by the SEM. The SEM micrograph analysis indicated that the mean nanoparticle diameter is 20.5±5.2 nm (**Figure 1 a**). The TEM film cross-section analysis of the FIB cut lamella revealed that nanoparticles are distributed over the whole 100 nm thickness of the film and are *ca.* 2 times smaller than nanoparticles visible on the surface (**Figure 1 b, d, f, h**). Some layering is seen at the initial stage of the film growth that was earlier observed experimentally by Jurkevičiūtė *et al.* ^54^ and analysed theoretically by Kairaitis *et al.*^55^ where it is explained as a dominating layered growth mechanism over columnar one at lower substrate temperatures due to the lower diffusion coefficient.

**Figure 1.**
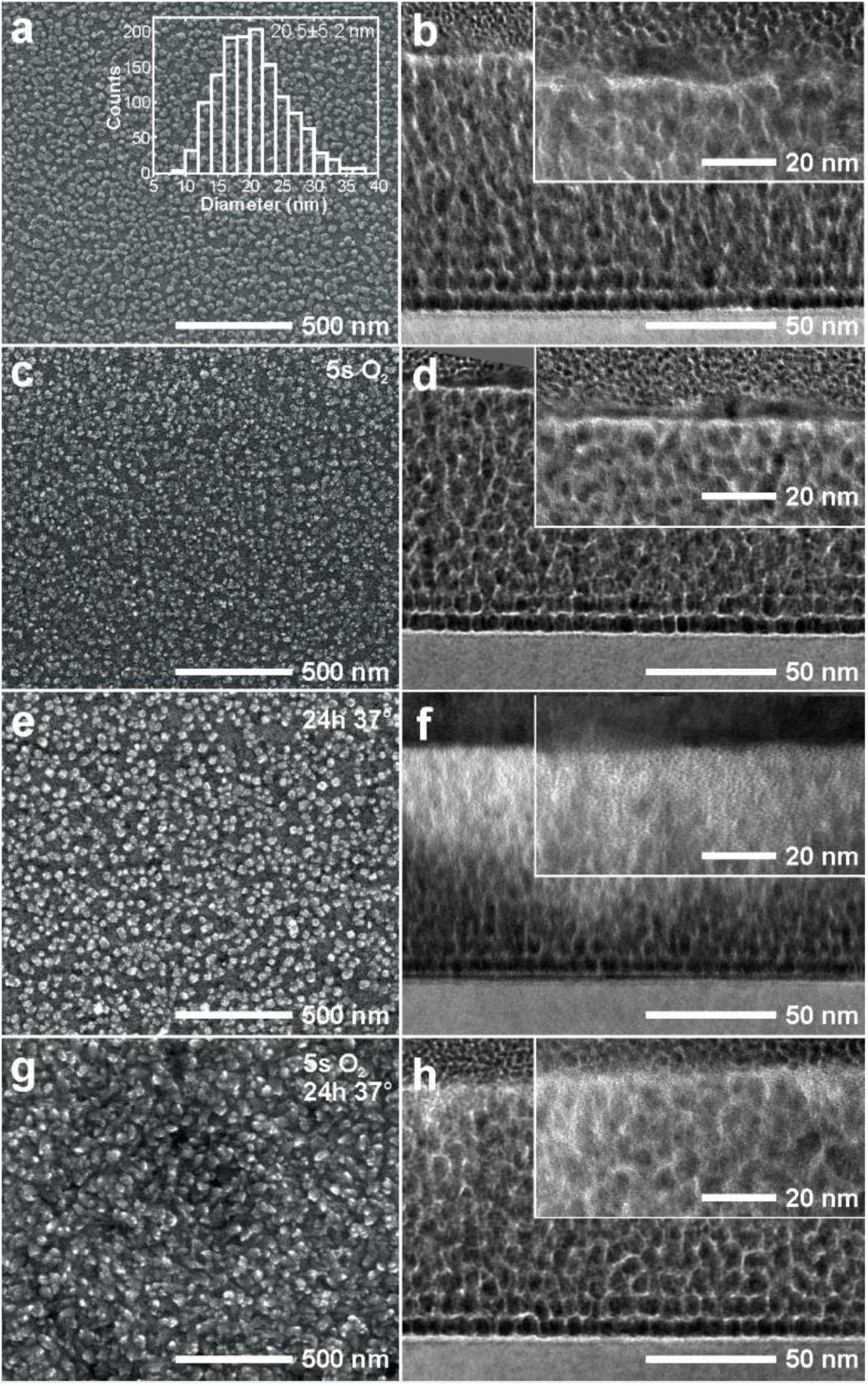
DLC:Cu film surface SEM (a, c, e, g) and cross-section TEM (b, d, f, h) micrographs of pristine film (a, b), 5 s O_2_ plasma processed (c, d), immersed in pure water for 24 hours at 37°C (e, f), and affected by both, plasma processed and exposed to water, with the identified length scale bars. The inset in (a) depicts nanoparticle diameter distribution. The insets in TEM micrographs depict a higher magnification view of the area close to the surface. The uniform colour area in (d) is due to the rotation of the original micrograph.

The SAED patterns of the TEM lamella depicted in (**Figure. 2 a)** were addressed to Cu (111), (002), and (022) which confirms presence of metallic Cu. The presence of copper Ⅱ oxide in the films was confirmed electron diffraction rings addressed to (011), (112), and an (224). High resolution TEM images of the same sample also indicated presence of metallic and oxidized copper in Cu_2_O state as depicted in (**Figure 2 b**) where 0.195 nm and 0.290 nm atomic plane distances were identified. The latter phases were also found in the grazing angle XRD diffractograms (**Figure 2 d**). The positions of the diffraction peaks are consistent with the PDF 004-0836 and 074-1230 for crytaline Cu and Cu_2_O phases, respectively.

**Figure 2.**
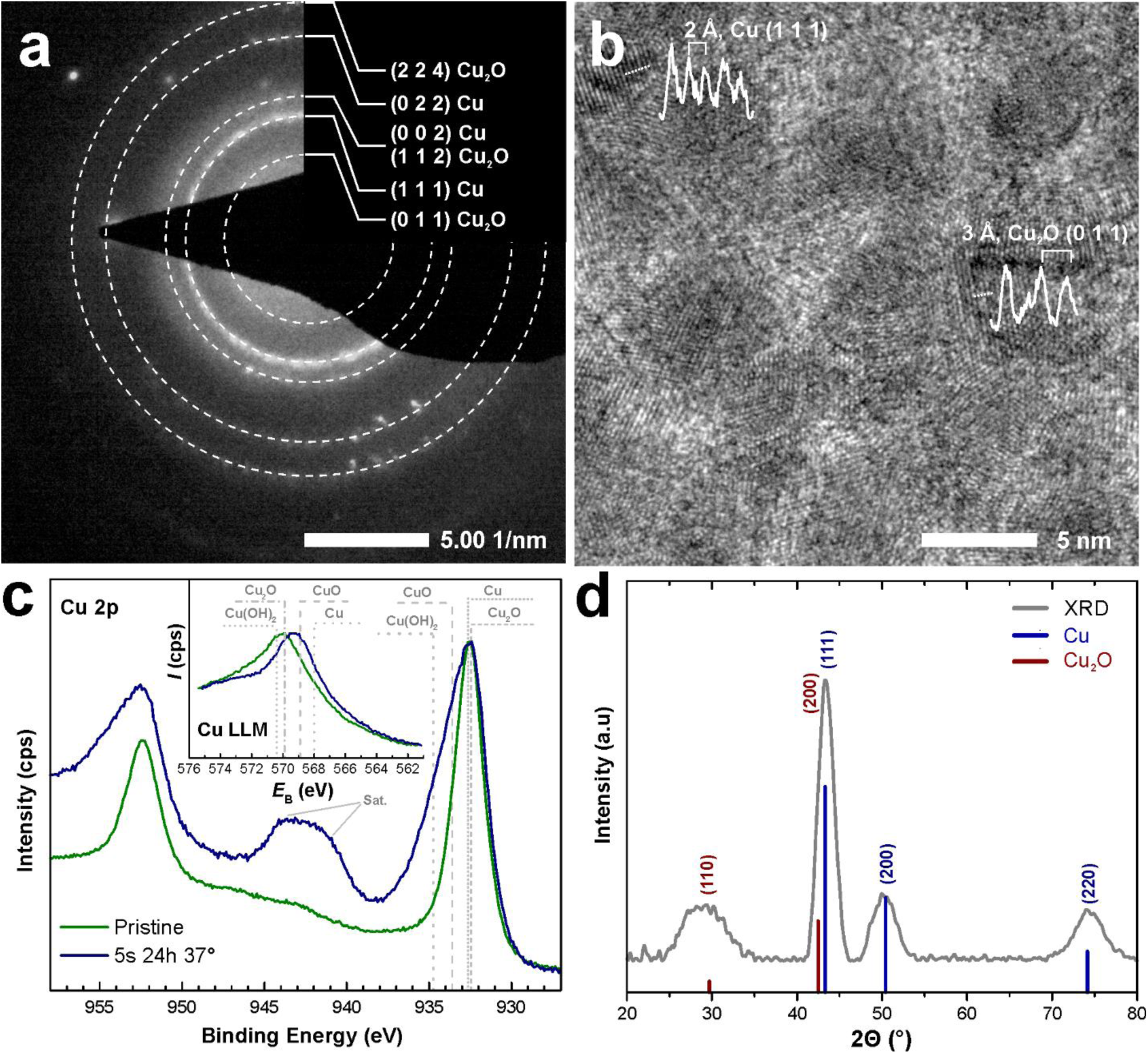
Chemical and structural analysis of DLC:Cu film. (a) SAED of the film lamella with identified *hkl* indexes and phases. (b) TEM lamella of the films with facets addressed to Cu and Cu_2_O, (c) XPS Cu 2p region spectra of pristine and 5 s O2 plasma treated and 24 h water exposure at 37C DLC:Cu film. Inset depicts Cu LMM region spectra. Differently dashed lines indicate energy values for known copper bonds. (d) XRD patterns of the DLC:Cu film with *hkl* indexes.

The XPS spectra in Cu 2p region of pristine DLC:Cu films showed two high-intensity peaks at 932.5 eV and 952.2 eV (**Figure 2 c**) corresponding to copper Cu 2p_3/2_ and Cu 2p_1/2_ doublet, respectively. There, the peak at 932.5 eV could be assigned to copper in Cu_2_O since its position is in good agreement with reported^56, 57^ (932.45 eV, 932.43 eV) binding energy values for cuprous oxide. A similar binding energy value is reported for metallic copper (932.607 eV, 932.61 eV, and 932.63 eV)^56–58^, therefore the presence of metallic copper cannot be excluded based on this peak position. Low-intensity satellite peaks at approximately 943 eV and 947 eV correspond to metallic copper or copper in Cu_2_O^57^. A similar main peak position was found for the rest of the samples in the Cu 2p region (**Figure S1**).

To investigate the oxidation state of the copper, representative Auger spectra in Cu LMM region for the same DLC:Cu samples was acquired (inset **Figure 2 c**). For the untreated film the main peak is positioned at 569.9 eV and coincides with known binding energy values for Cu_2_O (569.89 eV, 569.61 eV)^56, 59^, indicating presence of Cu(I) copper in the form of cuprous oxide (Cu_2_O).

The pristine DLC:Cu films withstood the surface abrasion test. The impact on the surface after different mechanical treatment conditions are depicted in **Figure S2**. The number and area of scratches increased with the decreasing sandpaper grits and accumulated contact and rotation speed.

### 3.2 Properties of the treated DLC:Cu films

For the antimicrobial analysis the experiments are usually carried out in aqueous environments mimicking the natural environment and human temperature therefore it is important to investigate their behaviour in similar time frames and temperature ranges ^12^. Moreover, as was reported earlier for Ag nanoparticles containing nanocomposite films, they released metal ions more effectively after short O_2_ plasma processing ^27, 37^. The effect of short plasma processing duration appears as contrast at the nanoparticle edges in the SEM micrographs (**Figure 1 c**), while in the TEM analysis, no obvious differences are seen (**Figure 1 d**). Pristine film immersion in ultra-pure water results in well-expressed contrast change over the film thickness (**Figure 1 f**) that could be attributed to the decrease in Cu content compared to the lightweight amorphous DLC matrix. Plasma-processed films seem to release Cu only from the very top layers (**Figure 1 h**) although their surface morphology became more pronounced after exposure to water (**Figure 1 g**).

Elemental analysis obtained with EDS indicated that plasma processing increases the oxygen content in the films and decreases the relative amount of Cu (**Figure 3 a**, **Table S2**) but changes are minor. Immersion in water makes a bigger change in decreasing copper content whereas in combination with plasma pre-processing immersion makes an extra effect on the increased oxygen content.

**Figure 3.**
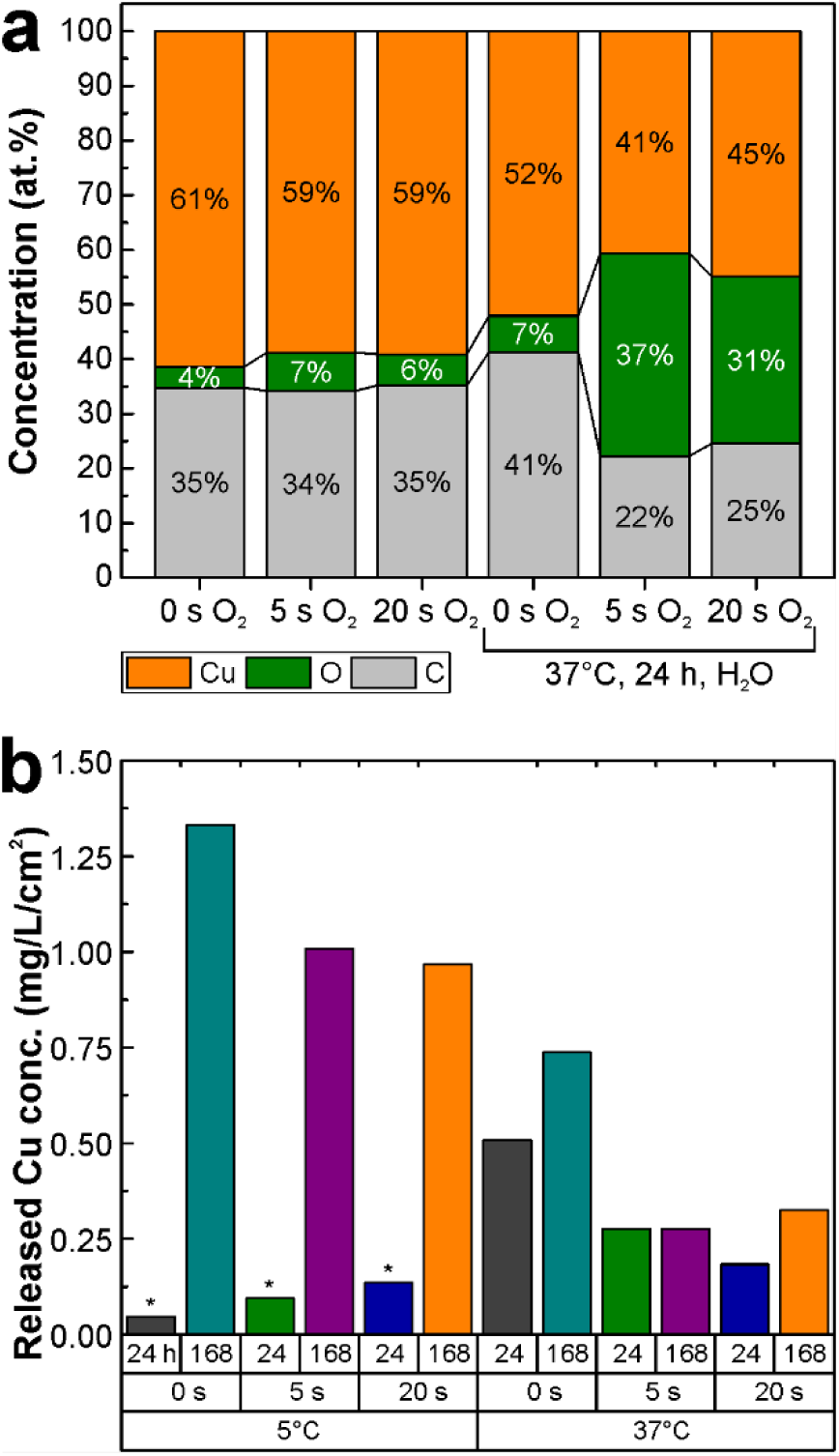
Pristine DLC:Cu and O_2_ plasma processed film elemental composition in atomic percent obtained by EDS before and after exposure to ultra-pure water for 24 h at 37°C (a) and AAS measured copper content released in ultra-pure water after plasma processing and immersion exposure condition indicated below the histogram (b). Asterix indicates results close to the measurement detection limit.

Plasma and water exposed DLC:Cu sample XPS analysis of the copper Auger spectra in Cu LMM region (inset **Figure 2, c**) indicated main peak at 932.5 eV which overlaps with the peak for the pristine DLC:Cu film, but is much wider and demonstrates clear satellite peaks at 941 eV and 944 eV, clearly indicating that Cu(II) is present in Cu(OH)_2_ or/and CuO bonds, as explained by Biesinger *et al.*^57^ and by Midander *et al.*^33^. The XPS spectra in Cu 2p region for investigated samples showed spectra similar to the pristine DLC:Cu film (**Figure S1 a**) except for sample immersed in virus solution. The latter sample also demonstrates low-intensity satellite peaks typical for divalent copper (at 941 eV and 944 eV).

The same assumption is also true for the sample treated in oxygen plasma for 5 s, for sample immersed in water for 24 hours and sample immersed in virus solution (**Figure S1 c**). On the contrary, in the spectrum for the sample treated for 5 s in oxygen plasma and immersed in water for 24 h at 37°C (**Figure 2, b**) the main peak at 569.3 eV corresponds to known CuO position (568.72 eV, 569.03 eV)^56, 59^ and indicates presence of Cu(II) in CuO form. This peak also shifted towards Cu_2_O position, suggesting presence of small amount of cuprous oxide. It should be mentioned, that the sample immersed in virus solution (no plasma treatment) (**Figure S1 c**) showed presence of Cu(OH)_2_ bonds (570.4 eV, 570.35 eV)^56, 59^. This assumption is in agreement with low-intensity satellite peaks typical for divalent copper found in Cu 2p region (**Figure 2**, a), discussed in the section 3.1 for this sample. Neither of all these samples exhibited a peak at 568 eV for Cu(0), this suggests a lack of metallic copper on the surface of the films, as is described in more detailed in the literature^59, 60^ or possible metallic copper core inside the Cu_2_O shell^33^.

XPS spectra in O 1s region (**Figure S1 c**) also showed different spectra for plasma treated sample and for plasma treated sample and immersed in water. There the presence of Cu(ll) in both CuO and Cu(OH)_2_ bonds can be assumed, although the C=O, C-O-C bonds positions and Cu(OH)_2_ positions are too close to reliably state that the Cu(OH)_2_ bonds are present^56^. As was demonstrated in Cu LMM region, the presence of Cu(OH)_2_ is unlikely (or present in small quantity), thus we conclude, that for this sample (**Figure S1 c**) the copper atoms are present in Cu_2_O and CuO bonds. Overall, the analysis O 1s region results supports findings in Cu 2p and Cu LMM regions. In contrary to O 1s region, all samples demonstrated almost identical carbon C 1s spectra (**Figure S1 d**), showing that most of carbon atoms are present in C-C bonds (sp^3^ hybridization), as expected for DLC films^60^. The small amount of carbon in C=C bonds (sp^2^) and in carbon-oxygen (C-O, C=O, O=C-O) bonds is also present.

The changes seen in the electron microscope micrographs are in line with the released Cu compound concentrations in ultra-pure water where after 24 h exposure at 37°C pristine sample released more metal content than 5 s plasma processed one (**Figure 3 b, Table S3**). In most of the cases, longer immersions resulted in a bigger metal release. Longer plasma processing did not increase the Cu release as was initially expected.

Oxygen plasma processed DLC:Cu sample visually appears almost identical to pristine samples, *i.e.*, copper red, but after exposure to water they change their colour into green shades characteristic of colored surface films (patina) that form on copper alloys ^61^ (**Figure S3**). These color changes are related to copper oxidation to cuprous oxide in air and further reaction into cupric oxide that in the presence of carbon dioxide and humidity turns into copper carbonate hydroxide or otherwise malachite ^62–65^.

As seen from the SEM micrographs (**Figure 1**) both O_2_ plasma processing and exposure to water modify the surface therefore the roughness of the samples was also inspected with AFM, and it was confirmed that pristine samples have lower average and root mean square roughness than water exposed samples (**Figure 4**). Immersion duration has a bigger effect on the increasing roughness than the durations of plasma processing.

**Figure 4.**
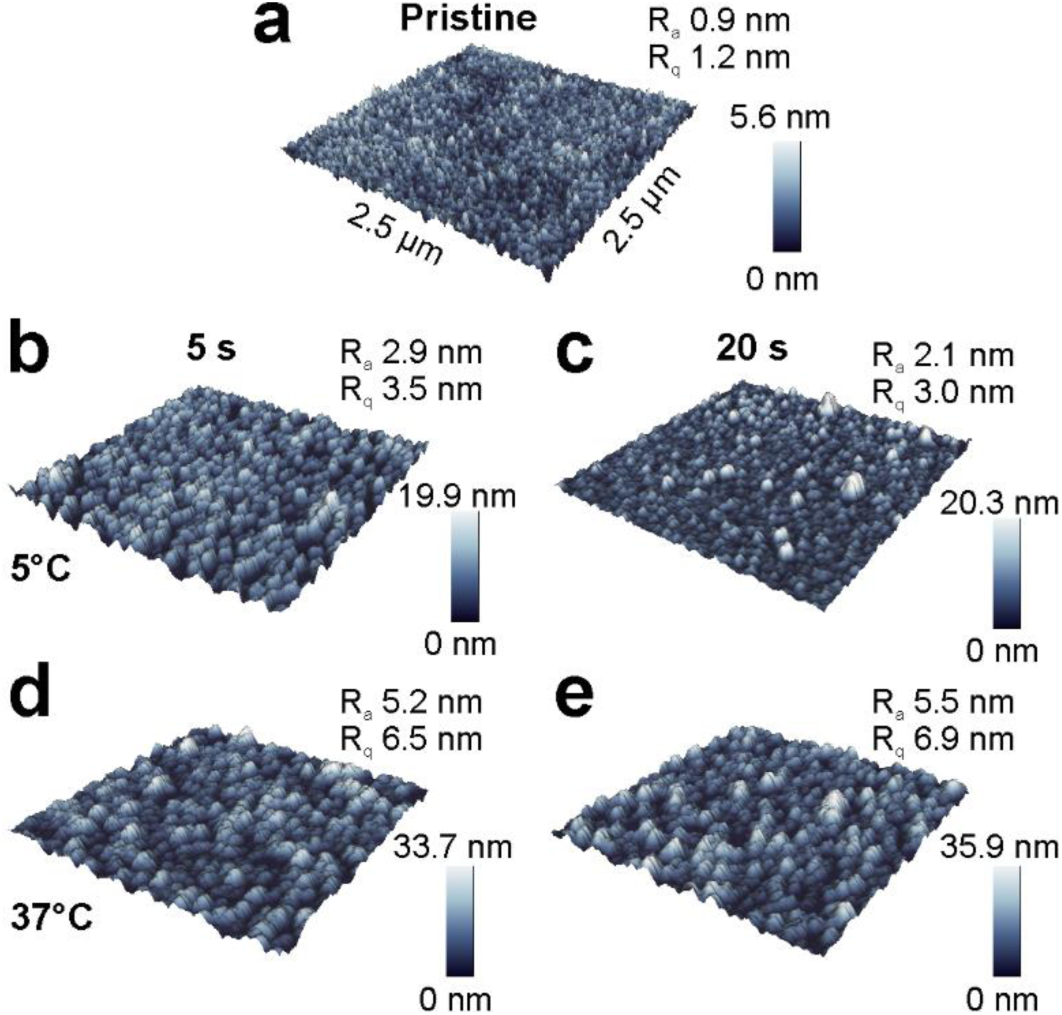
Pristine (a) and ultra-pure water exposed DLC:Cu film 2.5 µm x 2.5 µm sized area topographies obtained with AFM depending on the plasma processing durations 5 s (b, d) and 20 s (c, e) and immersion temperatures 5°C (b, c) and 37°C (d, e). Samples were immersed for 168 h. Numerical values in the top right corner indicate average roughness (R_a_) and root mean square roughness (R_q_).

After exposure to water changes in the nanocomposite films resulted in the changes not only in the loss in metal filler but also in some slight modification of the DLC matrix. As seen from the Raman spectra (**Figure 5 a, b, c**) and their deconvolution parameter using two typical D and G bands representing *sp*^2^ and *sp*^3^ bonded carbon phases (**Figure 5 d, e, f**) the DLC matrix is stable. The oxygen plasma treatment does not influence the G peak position (1546-1547 cm^-1^). It was previously observed that DLC film processing by O_2_ plasma can increase the *sp*^3^*/sp*^2^ ratio ^66^. Compared to pristine film the G peak tends to redshift by a maximum of 15 cm^-1^ and it is more prominent under higher immersion temperatures (**Figure 5 d**). According to the 3-stage model described by J. Robertson ^67^, the initial G peak position is in Stage 3 which shows the evolution of the structure from a-C carbon to ta-C carbon. The shift of G peak position is related to the increase of *sp*^3^ amount in the films. The G peak FWHM is a measure of disorder and increases with an increase of disorder ^68^. The biggest changes in the peak width are observed at higher temperatures but the G_FWHM_ varies only by 12 cm^-1^ (**Figure 5 e**). The variation is uneven as at shorter periods (24h) and lower temperatures (5°C) the G_FWHM_ slightly increases, whereas keeping in water for longer periods decreases the values of G_FWHM_. When samples were kept at a higher temperature in water, the variation of G_FWHM_ is more expressed which can be related to the rearrangement of surface layers as more oxygen is incorporated into the surface and less copper is released into the water. The peak intensity ratio I_D_/I_G_ varies depending on the applied treatment. Tendencies are changing depending on the applied temperature. For low temperatures independently of plasma treatment, the I_D_/I_G_ ratio is increasing indicating decrease of *sp*^3^ fraction in the films ^68–70^ which is contraversary to the redshift of G peak position. The intensity ratio fluctuates more when samples are kept at higher temperatures. The variation in I_D_/I_G_ ratio (0.45 – 0.65) could be attributed to the signal enhancement of copper NPs observed on the surface after oxygen plasma treatment. Copper incorporation into DLC affects its structure and a higher concentration of Cu reveals in *sp*^2^ richer structure ^14^. The observed minimal changes in the peak shape (**Figure 5 a-c**) and revealed parameters like G peak position, FWHM, and I_D_/I_G_ ratio support the idea that despite microscopic and visual changes of the films which are mainly related to Cu oxidation the DLC matrix stays almost intact with minor structural change compared to pristine DLC:Cu films.

**Figure 5.**
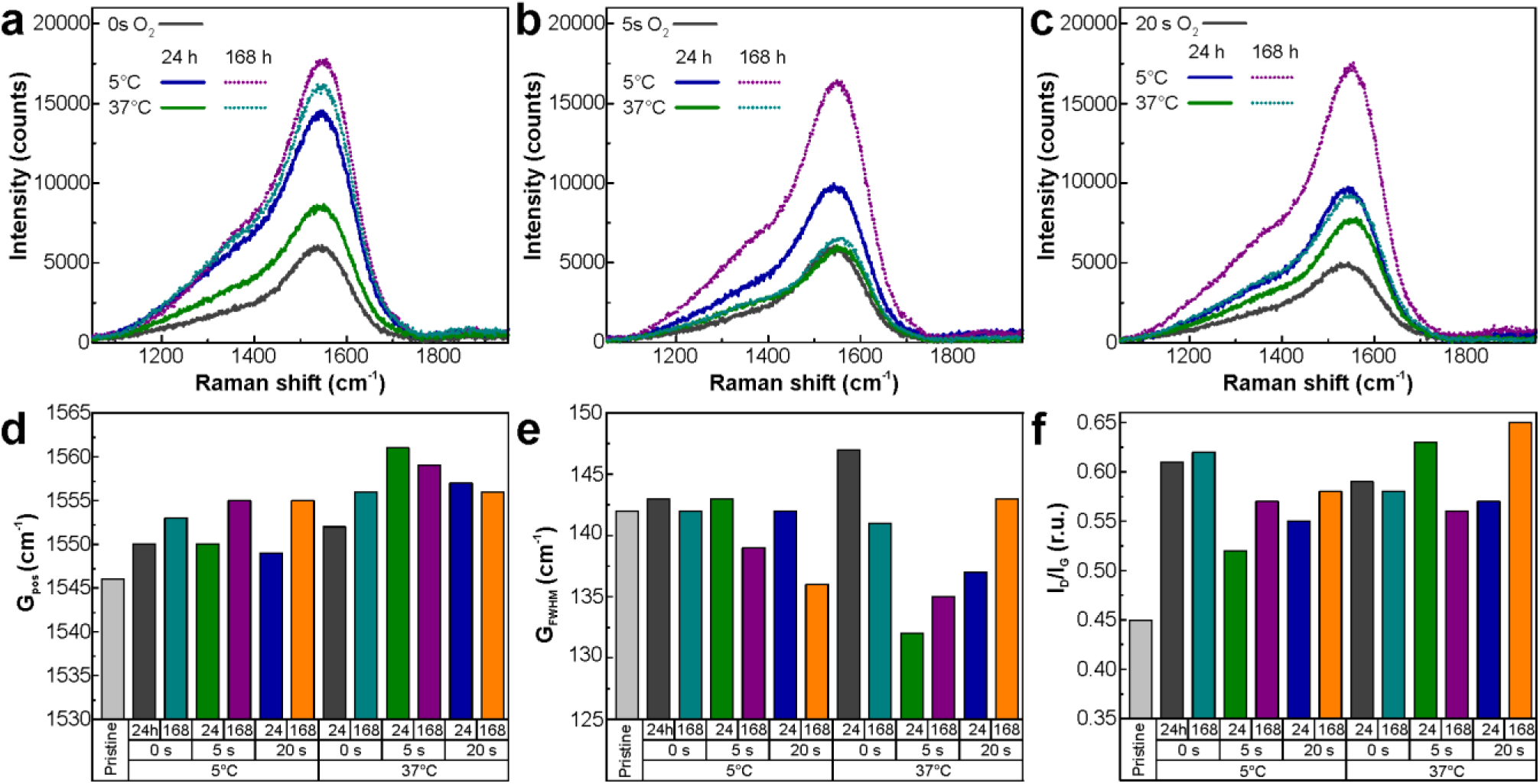
DLC matrix structure Raman analysis of pristine (a), 5 s (b), and 20 s (c) oxygen plasma processed DLC:Cu films after exposure in ultra-pure water at different conditions as indicated in the legends. The deconvoluted spectra into *sp*^2^ and *sp*^3^ phases expressed as G position (G_pos_) (d) and G peak full width at half maximum (G_FWHM_) (e), G and D peak intensity ratio (I_D_/I_G_) (f) after different sample exposure indicated below the histograms.

### 3.3 Cytotoxicity test

Cytotoxicity test results for MDBK and Vero cells provided in Tables **S4** and **S5**, respectively, and depicted in **Figure 6** indicate that the 100% extract obtained by incubating the cell culture media with 5% FBS at 37°C for 24 hours is cytotoxic. The cytotoxic properties gradually decrease with the titration of the extract. A decrease in cell viability was confirmed by the MTT assay where the increase of number of dead cells and a decrease in the metabolic activity and/or number of live cells was measured by the change in optical density (570 nm) compared to the control. The test showed a decrease in the metabolic activity of MDBK cells up to a 1:16 dilution (6.3%) and Vero down to a 1:8 dilution (12.5%). Metabolic activity was less than 50% for MDBK at 1:8 dilution and for Vero at 1:2 dilution. CC_50_ calculations showed that a 50% decrease in viability of MDBK is caused by 10.1% of the treated medium, while Vero - 30.0%.

**Figure 6.**
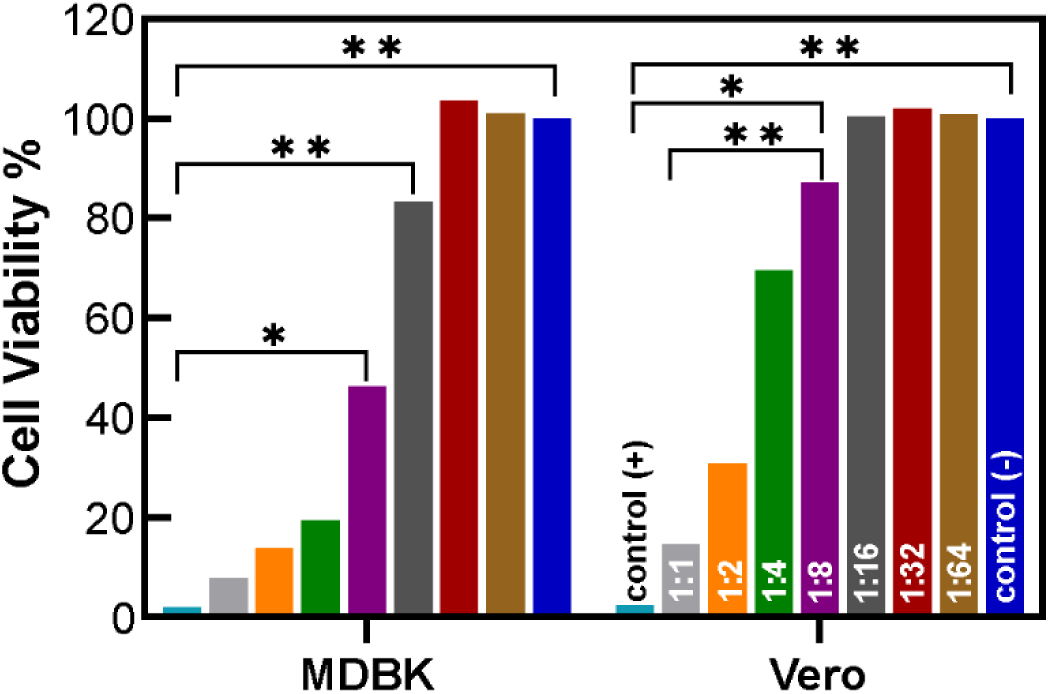
MTT assay results showing cell viability under different DLC:Cu film extract titration over 24 h. (\**p* < 0.04, \*\**p* < 0.001).

Inspected cell morphological changes are seen in **Figure S4**. Lysis of the MDBK cells was induced by exposure to an undiluted medium and diluted by two-fold dilutions up to 1:32 (3.2%). Undiluted and 50% medium caused nearly a complete destruction of cell layers (grade 4). Less cytotoxic was the 25% medium, which caused changes or lysis of about 50-60% of the MDBK monolayer cells, but not complete destruction of the monolayer, as is characteristic of grade 3. With further dilution, the cytotoxicity decreased and the 12.5% medium caused up to 50% of the cell lesions, and cell lysis was rarely observed as characteristic of grade 2. Dilutions with 6.3% and 3.2% medium caused up to 5- 10% cell lesions, and single cell lysis was observed as characteristic of grade 1. While for Vero cell morphology changes were induced by exposure to media up to a 1:16 (6.3%) dilution. Undiluted medium caused nearly complete destruction of cell layers (grade 4). After twofold dilution (50% medium) the cytotoxicity decreased and 50-60% change in cell morphology or lysis (disintegration) was observed, but no complete destruction of monolayer was observed, as is characteristic of grade 3. The cytotoxicity decreased further with dilution and 25% medium caused up to 50% cell changes, and cell lysis was observed rarely in grade 2. Low cytotoxicity was observed with 12.5% and 6.3% medium. These media caused up to 5-10% cell changes and single cell lysis, as is characteristic of grade 1. Slight alkalization of the medium was observed (**Tables S4** and **S5**).

### 3.4 Virucidal efficacy

DLC:Cu and DLC:Cu+O_2_ samples showed virucidal effects against both model viruses (**Figure 7**). In the case of IBV as depicted in **Figure 7 a** and **Table S6** DLC:Cu and DLC:Cu+O_2_ samples showed the same amount of decrease in the number of coronaviruses by 2.23 log_10_/ml TCID_50_/ml compared to the IBV control or 99.41%. The decrease in BoHV-1 titre was 1.63 log_10_/ml TCID_50_ or 97.7% for DLC:Cu sample, and 0.80 log_10_/ml TCID_50_ or 84.2% for the DLC:Cu+O_2_ samples compared to BoHV-1 control (**Figure 7 a** and **Table S7**). For the Si control sample, the titre of IBV and BoHV-1 decreased by 1.77 log_10_/ml TCID_50_ and 0.13 log_10_/ml TCID_50_, respectively.

**Figure 7.**
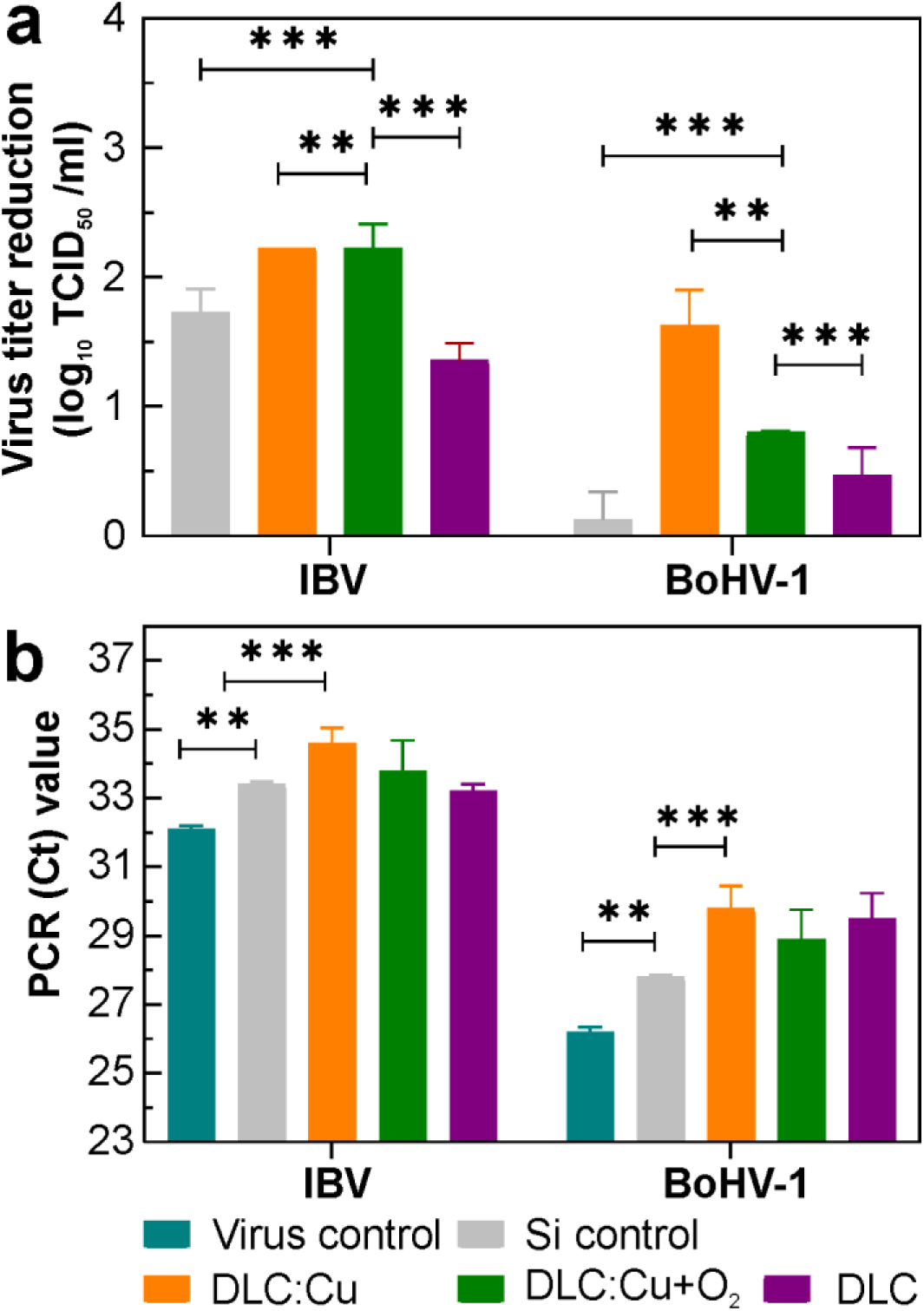
Antiviral efficacy of DLC, DLC:Cu, DLC:Cu+O_2_ films and controls, (a) virus titre log_10_ reduction of IBV, and BoHV-1 after 1 h of contact with films (\*\**p* = 0.004 and \*\*\**p* ≤ 0.0001). (b) RT-PCR cycle threshold (Ct) values (\*\**p* = 0.001 and \*\*\**p* ≤ 0.0001).

### 3.5 Real-time PCR test

The comparative analysis of IBV and BoHV-1 inactivation and viral bioactivity properties of the investigated samples by evaluating post-contact quantification of TCID_50_ titre (**Figure 7 a**) and RT-PCR threshold cycle Ct values (**Figure 7 b**) showed similar tendencies. The PCR Ct values for IBV ranged from 33.2 for DLC to 34.6 for DLC:Cu (**Figure 7 b, Table S6)**. The investigation of DLC:Cu sample showed statistically reliable (*p*<0.0001) rates of IBV RNA degradation and directly influenced the PCR Ct value. While Ct for the BoHV-1 ranged from 28.9 for DLC:Cu+O_2_ (*p*<0.001) to 29.8 (*p*<0.0001) for DLC:Cu (**Figure 7 b, Table S7**). The increase in Ct value can be attributed to virus inactivation properties of DLC:Cu sample, which passively reduces the number of viruses in control samples when they come into contact with virus particles.

For a better impression of the Cu release from the DLC:Cu in the virucidal study, the preliminary analysis of the aqueous medium volume and thickness above the immersed sample influence on the released Cu concentration was conducted in DMEM. Two different film area samples, namely 2.1 cm^2^ and 55.1 cm^2^ were immersed in the same 5 ml volume of liquid and different size Petri dishes.

The AAS measurement results depicted in **Figure S4** indicated that accumulated Cu concentration is bigger and approaching 2.5 mg/L/cm^2^ when the liquid volume was relatively big compared to the area. For the big area samples, where aqueous medium was just covering the sample surface the Cu release was almost 10 times smaller in the same solvent volume. If compared with the Cu release in ultra-pure water (**Figure 3b, Table S3**) the release in DMEM is faster. And that might contribute to positive virucidal and antibacterial results.

### 3.6 Virucidal Efficacy Stability

The testing of coating virucidal durability properties (**Figure 8**) indicated a gradual decline of virucidal activity against BoHV-1 from 3.00 log_10_ to 1.05 log_10_. The best virucidal effect was achieved by the pristine sample, which reduced the number of viruses by 99.9% within 1 hour, *i.e.*, from 7.25 log_10_ to 4.25 log_10_. After this, film was kept additionally with cell culture medium for 2 hours, it was used for the second virucidal test and reduced the number of viruses by 99.5%. The third and fourth cycles of testing of the virucidal effect of the coating showed that more than 90% BoHV-1 were inactivated (see **Table S8**).

**Figure 8.**
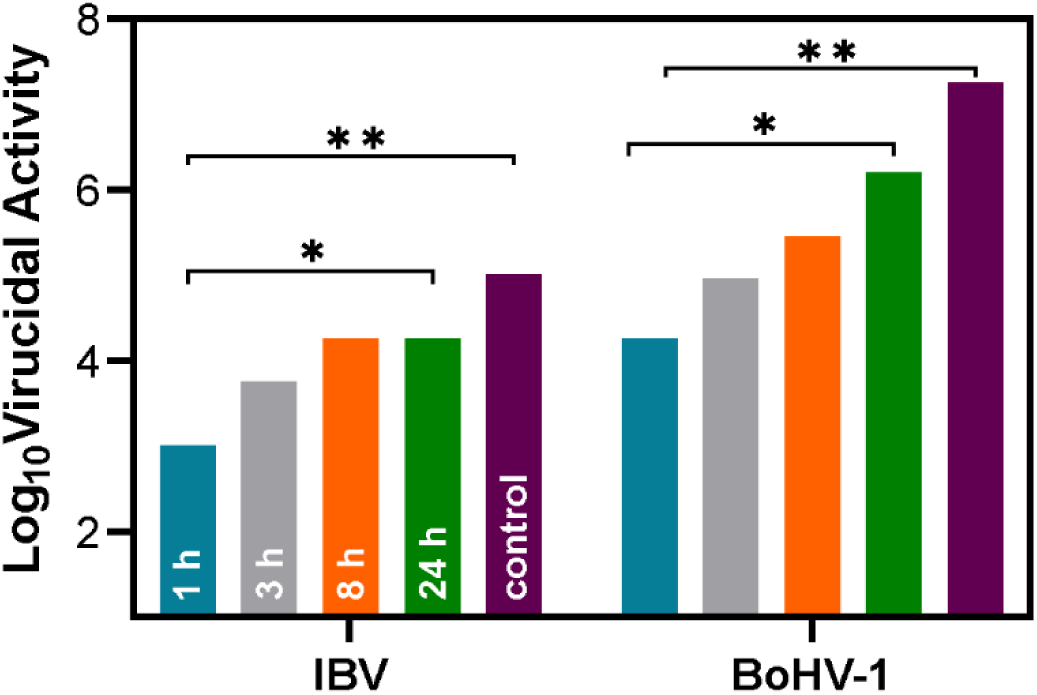
Evolution of virucidal activity of the DLC:Cu film after 1, 3, 8, and 24 h (\**p* < 0.045, \*\**p* < 0.009).

As for IBV the virucidal activity gradually decreased from 2.00 log_10_ to 0.75 log_10_ during coating durability test. The best virucidal effect was obtained for pristine film, which reduced the number of viruses by 99.0% within 1 hour, *i.e.*, from 5.0 log_10_ to 3.0 log_10_. In the second inspection the number of viruses was reduced by 94.38%. The third and fourth virucidal tests of the coating showed that 82.22% of IBV was inactivated after both 8 and 24 hours of exposure (**Table S9**).

### 3.7 Antibacterial properties

As the plasma pre-processed DLC:Cu films exhibited a lower virucidal effect than the pristine DLC:Cu films the antibacterial study was limited only to the latter. The antibacterial study showed that the DLC:Cu films had an expressed negative effect on the survival of both investigated bacteria strains. The results were confirmed by the identical tests which gave very similar results. No gram-negative *E. coli* bacteria were found in all samples after 3 hours as well as over longer periods, resulting in 100% antibacterial efficiency (**Figure 9 a, Tables S10-S12**).

**Figure 9.**
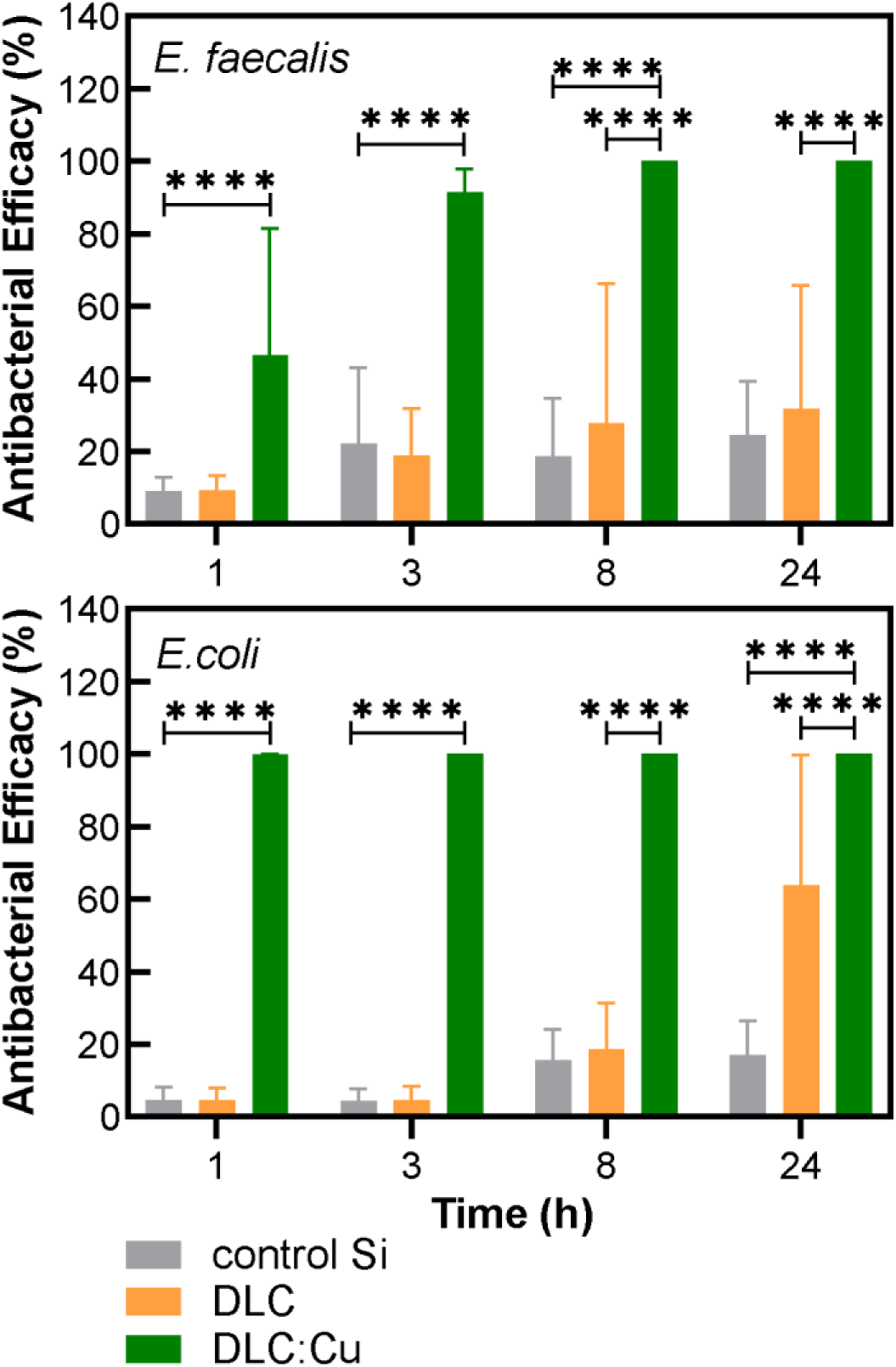
Antibacterial efficacy of the DLC:Cu obtained after immersion in (a) *E. coli* and (b) *E. faecalis* PBS broth and sampling after different periods from 1 to 24 hours. The impact of silicon substrate and DLC alone is indicated for reference (\*\*\*\**p* ≤ 0.0001)

The study employing gram-positive *E. faecalis* bacteria after 3 hours in Test 1 resulted in 86.87 ±1.7 % antibacterial efficacy and bacteria-killing half-life of 61.9 ±3.8 min (**Figure 9, Table S13**) and in Test 2 resulted in 99.97±0.01% and 15.2 ±0.18 min (**Table S14**) and in test three resulted in 89.300±0.42 and 60.63±1.9 min (Table S15). After longer investigated durations no *E. faecalis* bacteria were found in all samples and 100% antibacterial efficiency was obtained.

The silicon substrates used as a reference had a smaller effect on tested bacteria and a decrease in the number of bacteria was more pronounced over longer testing periods (**Figure 9 b, Tables S13- S15**). Images and the log_10_ CFU/ml graphs of both investigated bacteria shows the effect of the DLC:Cu films after 24 hours, compared to Si and DLC without Cu, the images shows that DLC without Cu have lower effect on both bacteria, while the DLC:Cu films results in 100% bacteria loss (**Figure S5**).

## 4. DISCUSSION

The electron and atomic force microscopies confirmed the changes in the DLC:Cu films surface after exposure to ultra-pure water depending on the plasma pre-processing that was primarily recognized as changes in colour seen by the naked eye. The Cu nanoparticle filler tends to be released from the pristine samples faster than compared to the plasma-processed ones and over 24 h it is released more effectively at human temperature than in a cool environment. These differences could be beneficial where controlled release over longer periods is expected and, in such cases, short surface modification with O_2_ plasma could prolong the use of nanocomposite DLC:Cu films. Over a longer period of 168 h more Cu is released in the cool environment because probably it is already saturated over the same period at 37°C. The concentration range of Cu released in water was up to 1 mg/L/cm^2^ and it is at a similar level as it was reported for carbonaceous matrix nanocomposite films with silver where Juknius *et al.* ^27^ measured up to 0.4 mg/L/cm^2^ Ag release from O_2_ plasma processed DLC:Ag film after 2-day soaking in 35°C 10 ml water, while Merker *et al.* ^12^ obtained up to 0.1 mg/L/cm^2^ of Ag in 20 ml water at 7-days immersion at 37°C depending on the protective ultra nanocrystalline diamond film thickness. The study of Tamayo *et al.* ^28^ covered longer time immersion periods in 10 ml of water extending over 3 months and resulting in Ag release of 0.35 mg/L/cm^2^ for polyethylene nanocomposites with Ag nanoparticles and 90 mg/L/cm^2^ with Cu. A big difference between Ag and Cu was addressed by the fact that Cu is susceptible to oxidation. The latter was confirmed by the XPS analysis where for pristine DLC:Cu films copper is mostly present in the form of cuprous oxide (Cu_2_O). While after immersing in the water, a plasma treated sample resulted in the appearance of Cu(II) in the CuO form. Also, after virus treatment of pristine sample it resulted in appearance of small amount of Cu(II) in the Cu(OH)_2_ form.

Differences in the Cu compound release in the case of plasma-processed samples indicate different surface properties of oxidised Cu. This might be related to differences in solubility as reported by Wei *et al.* ^71^ where Cu salt fillers tended to release more Cu than metallic filler. Based on the elemental changes after water exposure detected by EDS and changes in sample colour, it suggests that oxidation and maybe hydroxide formation took place ^61^. Dissolution and depletion of metallic NPs were explained theoretically by Quezada *et al.* ^72^ employing a shrinking core model where it is assumed that while the NP reacts with the environment it forms a soluble product and a porous layer called shell, which is formed around a solid core of the NPs.

The actual virucidal and bactericidal testing experiments are usually performed not in ultra-pure water but in different media. Therefore, Cu ion release experiments were reported previously in a range of solvents, for example, Wu *et al.* ^29^ investigated Cu release from mesoporous silica xerogels in standard simulated body fluid, and the released concentration correlated with the initial Cu content. It was also reported that alloyed composite implants tend to release their compounds differently depending on the solvent pH ^30, 31^. In our study, a bigger Cu concentration registered by AAS after immersion in DMEM containing 2.0% of bovine fetal serum compared to ultra-pure water could be attributed to the influence of the different aqueous medium composition and therefore differences in the surface effects. The observed saturation of released Cu over time is related to the volume of the solvent and concentration gradient in the liquid layer. The observed logarithmic release trends were reported by others as well ^29, 35, 71, 72^ and could be related to limited-source diffusion in the case of thin films with low total metal volume fraction. This preliminary study of short-duration immersion in DMEM confirmed that Cu release from DLC:Cu film is not momentarily, and it also suggests that a significant amount of Cu should have been released during the 1-hour antiviral testing. Nevertheless, having in mind a small aqueous medium gap in the actual experiment, *c.a.* 100 µm and a small volume (10 µl) the Cu release might be limited similarly to the big-sample-thin-layer example. While in the antimicrobial testing, where samples were immersed in PBS broth Cu release should have been more pronounced. This hypothesis cannot be directly confirmed with the AAS measurements because it requires bigger sampling volumes than available in the actual virucidal testing experiments.

Summing up, the expected improvement in oxygen plasma processing observed for DLC:Ag films did not translate in the DLC:Cu study and the obtained differences could be addressed to the very different stability of the Ag and Cu metals and changes of the Cu compound release after oxidation. Most of the studies suggested that the generation of reactive oxygen species (ROS), protein oxidation, lipid peroxidation, and DNA degradation in bacteria are the major mechanisms of antibacterial action of the Cu NPs ^73^. The Cu ions released from surfaces was confirmed to have a direct action on a variety of targets and/or result in the production of ROS, leading to the nucleic acids denaturation and damaged virion integrity ^74^. Such mechanisms of action could also contribute to the inactivation of viruses. It was reported that during the incubation of the virus-containing droplet on investigated coatings, the virus particles may be inactivated both by their direct interaction with the surface and by their interaction with the released Cu ions. Upon consideration of the differences in sizes of the virus particle and the ions released in the liquid solution, their diffusion coefficients (Stokes–Einstein equation) and resulting transport speeds will be very different, where the virus particle migration is much slower than that of ions, indicating that any antiviral effect from the ions would occur faster ^75^.

Our studies showed that the cytotoxicity of copper-coated surfaces depended on the concentration of the tested sample. Therefore, it could be assumed that, the cytotoxicity of copper coatings is determined by the amount of copper entering the medium^76^. The amount and concentration of copper released from the coating into the medium can reach a maximum over a period of time depending on the amount and temperature of the incubated medium. The amount and dynamics of copper released from the coating differ (may vary) at the room and body temperature. It is likely that the maximum copper concentration will be reached more quickly when the medium is exposed to higher temperatures and its content is lower. The amount and concentration of copper released may not reach a maximum if the incubation or exposure is for a shorter period. In our studies at 37°C from the coating to the medium in 24 h. released copper showed cytotoxicity to MDBK and Vero cells after 24 h. incubation. The study showed that microscopic damage to single cells did not affect the total viability determination by the MTT test.

Durability studies of the virucidal effect of the copper coating were performed using four one-hour virus incubation cycles. Between these cycles, the coatings were exposed to cell culture media for 2, 4, and 15 h, during which the released copper reduced the virucidal potential of the coating. Our study showed that exposure of copper coatings to cell culture medium between cycles had an effect on virucidal activity^77^ against both BoHV-1 and IBV viruses.

The decrease in virus titre and quantity of nucleic acid detected after contact with DLC:Cu films were obtained. The differences in virus titre reduction after BoHV-1 treatment on DLC:Cu, DLC:Cu+O_2_, DLC, and Si control showed the key role of the Cu. The comparison of virucidal efficacy of DLC:Cu and DLC:Cu+O_2_ showed the importance of quicker Cu release and concentration leading to a difference of 0.83 log_10_ (up to 7 times) in virus reduction. The Cu influence in virucidal efficacy for BoHV-1 was also confirmed by RT-PCR. The real-time PCR detected a difference up to 2 times in the rate of DNA degradation after treatment on surfaces of pristine DLC:Cu compared to slower Cu-releasing plasma pre-processed films. The analysis of these results suggests that BoHV-1 proteins and nucleic acids are targeted by Cu nanoparticles or ions, but damaged at different speeds. BoHV-1 proteins located on the surface are probably targeted first and the DNA genome is affected later. A similar effect of Cu ions was detected by investigation of human herpes simplex virus DNA, envelope, and capsid biomolecules ^78^.

The reduction of virus titre after IBV treatment also indicated the key role of Cu.

The comparison of the virucidal efficacy of pristine DLC:Cu and plasma pre-processed film indicated similar virus reduction results despite the difference in the amount of released Cu indicated by the AAS analysis. The importance of the released Cu on the virucidal effect on IBV was confirmed by RT-PCR. The RT-PCR detected up to 1.5 times difference in the rate of RNA degradation after treatment on surfaces of pristine DLC:Cu compared to plasma-pre-processed film. Similarly like for herpes virus, the study for IBV suggests that proteins and nucleic acids are targeted by Cu nanoparticles or/and ions but might be damaged at different speeds. IBV protein capsid protects nucleic acid for replication, spike protein, membrane glycoprotein, envelope protein, and nucleocapsid proteins are located on the surface and are the first target while RNA genome sequences are affected later at significantly lower rates ^79–83^.

In general, a bigger decrease in viruses in the case of pristine DLC:Cu films compared to O_2_ plasma processed indicates the role of bigger released Cu content in the medium. Differences in the IBV and BoHV-1 titre reduction for investigated films imply that IBV is more dependent on the surface as a tremendous decrease was noticed on the film-free silicon surface. While BoHV-1, despite smaller virus reduction in absolute number was following the expected Cu release trend.

Bactericidal studies showed different kinetics of inactivation for bacteria possessing different types of cell walls. The tested bacteria, gram-negative *E coli,* and gram-positive *E. faecalis* are the most widely used indicators of faecal contamination ^84^. The main components of the *E. faecalis* cell wall are a thick layer of peptidoglycan, wall teichoic acids, and lipoteichoic acids, while *E. coli* cell wall has an inner and outer membrane, separated by a thin peptidoglycan layer. The different structures of bacterial cell walls can be a key factor in a different susceptibility or resistance to various factors and it might lead to the different kinetics of inactivation ^73, 85^.

*E. coli* bacteria were more sensitive (100% or 4.90 log_10_ reduction in 3 hours) to the bactericidal effect of DLC:Cu than *E. faecalis* but the hypothesis that investigated films are more effective against gram-negative bacteria would require a wider scope study ^86^.

Our research, evaluating the antiviral and antibacterial effectiveness of nanocomposite DLC:Cu films, demonstrated that surfaces containing copper nanoparticles showed a strong effect on both model viruses and bacteria.

## 5. CONCLUSIONS

Reactive magnetron-sputtered nanocomposite amorphous diamond-like carbon films with copper nanoparticles (DLC:Cu) effectively release the Cu compounds in the aqueous media and the total amount depends on the storing temperature, liquid film over the sample, and the medium itself.

Oxygen plasma pre-processing of the deposited DLC:Cu films oxidise the Cu and prolong the metal compound release in the medium which could be beneficial when controlled instead of immediate release is expected.

Presence of metallic copper and copper(I) oxide phases in the pristine DLC:Cu films were confirmed by multiple analytical methods, while XPS indicated presence of copper(II) oxide and copper(II) hydroxide especially in the differently exposed films.

The DLC matrix remains almost intact despite the different exposures to plasma and followed immersion in ultra-pure water as confirmed by the *sp*^2^/*sp*^3^ phase studies via Raman spectroscopy analysis.

DLC:Cu films were effective against both tested model RNA-containing coronavirus and DNA-containing herpesvirus as well as both tested gram-positive *E. faecalis* and gram-negative *E. coli* bacteria although differences in efficacy were observed.

The differences in the amount of released Cu from pristine and plasma-processed DLC:Cu films were directly seen in virucidal efficacy for both investigated viruses, but it was more expressed for BoHV-1 where it was observed as a decrease in virus titre and PCR cycle threshold value. While for IBV the released Cu concentration-related tendency was obtained only in Ct value.

Antibacterial study indicated that no *E.coli* were found after 3 hours of contact with pristine DLC:Cu films while for *E. faecalis* up to 99.97±0.25% antibacterial efficiency was obtained and 14.86±1.15 min bacteria killing half-life was determined. The differences could be addressed by the differences in the bacteria cell walls, but a wider range of bacteria should be tested to confirm the hypothesis.

## Supporting information

Graphical abstract

## ASSOCIATED CONTENT

### Supporting Information

- The equations used to calculate the half-life for the viruses and bacteria and antibacterial effectiveness.
- Pristine DLC:Cu and different duration O_2_ plasma processed film average elemental composition before and after exposure to ultra-pure water
- XPS spectra for different treated DLC:Cu samples.
- Optical microscope images of the DLC:Cu films after abrasion with different sandpaper grits, rotation per minute, and rotation cycles.
- AAS measured Cu concentration normalized to 5 ml solvent and 1 cm^2^ area released from pristine and different duration plasma processed DLC:Cu films in ultra-pure water incubated at different temperatures and durations.
- Camera images of 5 s and 20 s plasma processed DLC:Cu samples after immersion for 24 hours in 5°C and 37°C temperatures as indicated below the samples.
- Cytotoxicity results and optical microscope images of MDBK and Vero cells after incubation for 24 h in different dilution of extract obtained after DLC:Cu immersion in cell culture media. Virucidal testing results of IBV and BoHV-1 are summarized in Table S3 and Table S4 respectively.
- The antibacterial testing results that were repeated three times for each investigated *E. coli* and *E. faecalis* bacteria strains.
- Camera images of bacteria colonies grown on agar plates for Si, DLC, and DLC:Cu after 24 h immersion.

## Author Contributions

Conceptualization and methodology – T.T., A.Š., M. M.; Investigation – A.T., A.V., M.A., K.S., N.K., R.L., A.Š.; Data analysis N.K., M.M., R.L., A.Š., D.Z., S.B., A.T., T.T., Writing – original draft D.Z., S.B., M.M., N.K., A.T., M.A., T.T., S.T.; Writing – review & editing; D.Z., S.B., R.L., A.Š., Š.M., M.M., S.T., T.T., Supervision – T.T., A.Š., M.M.; Funding acquisition – T.T.

## Funding Sources

This project has received funding from the European Regional Development Fund (project No. 13.1.1-LMT-K-718-05-0018) under a grant agreement with the Research Council of Lithuania (LMTLT). Funded as the European Union’s measure in response to the COVID-19 pandemic.

## Notes

The authors declare no competing financial interest.

## ACKNOWLEDGMENT

A special thanks go to project No. 13.1.1-LMT-K-718-05-0018 members Dr. S. Račkauskas, Dr. R. Mardosaitė, M. A. Baba, G. Khatmy, M. Ilickas, Dr R. Zykus, Dr A. Urbas, M. Mikutis, O. Ulčinas, J. Baltrukonis, E. Nacius, M. Vainoris, D. Mazur from the Kaunas University of Technology, Lithuanian University of Health Sciences, and Altechna R&D for their technical assistance.

## Notes

### Competing Interest Statement

The authors have declared no competing interest.

### Summary of Updates

The manuscript was revised based on the reviewer's comments. New experiments were conducted and related new texts and figures (Figure 2, Figure 6, Figure 8, Figure 9) were provided in the main text and the supplements.

## REFERENCES

(1) Nwobodo, D. C.; Ugwu, M. C.; Anie, C. O.; Al-Ouqaili, M. T. S.; Ikem, J. C.; Chigozie, U. V.; Saki, M. Antibiotic resistance: The challenges and some emerging strategies for tackling a global menace. Journal of Clinical Laboratory Analysis 2022, 36 (9), 10, Review. DOI: 10.1002/jcla.24655.

(2) Saud, Z.; Richards, C. A. J.; Williams, G.; Stanton, R. J. Anti-viral organic coatings for high touch surfaces based on smart-release, Cu2+ containing pigments. Progress in Organic Coatings 2022, 172, 6, Article. DOI: 10.1016/j.porgcoat.2022.107135.

(3) Birkett, M.; Dover, L.; Cherian Lukose, C.; Wasy Zia, A.; Tambuwala, M. M.; Serrano-Aroca, Á. J. I. J. o. M. S. Recent advances in metal-based antimicrobial coatings for high-touch surfaces. 2022, 23 (3), 1162.

(4) Chang, T.; Sepati, M.; Herting, G.; Leygraf, C.; Rajarao, G. K.; Butina, K.; Richter-Dahlfors, A.; Blomberg, E.; Odnevall Wallinder, I. J. P. O. A novel methodology to study antimicrobial properties of high-touch surfaces used for indoor hygiene applications—A study on Cu metal. 2021, 16 (2), e0247081.

(5) Jastrzebski, K.; Bialecki, J.; Jastrzebska, A.; Kaczmarek, A.; Para, M.; Niedzielski, P.; Bociaga, D. Induced Biological Response in Contact with Ag-and Cu-Doped Carbon Coatings for Potential Orthopedic Applications. Materials 2021, 14 (8), 22, Article. DOI: 10.3390/ma14081861.

(6) Birkett, M.; Zia, A. W.; Devarajan, D. K.; Panayiotidis, S. M. I.; Joyce, T. J.; Tambuwala, M. M.; Serrano-Aroca, A. Multi-functional bioactive silver- and copper-doped diamond-like carbon coatings for medical implants. Acta Biomaterialia 2023, 167, 54–68, Review. DOI: 10.1016/j.actbio.2023.06.037.

(7) Sanzone, G.; Field, S.; Lee, D.; Liu, J. Z.; Ju, P. F.; Wang, M. S.; Navabpour, P.; Sun, H. L.; Yin, J. L.; Lievens, P. Antimicrobial and Aging Properties of Ag-, Ag/Cu-, and Ag Cluster-Doped Amorphous Carbon Coatings Produced by Magnetron Sputtering for Space Applications. Acs Applied Materials & Interfaces 2022, 14 (8), 10154–10166, Article. DOI: 10.1021/acsami.2c00263.

(8) Vincent, M.; Hartemann, P.; Engels-Deutsch, M. Antimicrobial applications of copper. International Journal of Hygiene and Environmental Health 2016, 219 (7), 585–591, Article; Proceedings Paper. DOI: 10.1016/j.ijheh.2016.06.003.

(9) Shigetoh, K.; Hirao, R.; Ishida, N. Durability and Surface Oxidation States of Antiviral Nano- Columnar Copper Thin Films. Acs Applied Materials & Interfaces 2023, 15 (16), 20398–20409, Article. DOI: 10.1021/acsami.3c01400.

(10) Meister, T. L.; Fortmann, J.; Breisch, M.; Sengstock, C.; Steinmann, E.; Koller, M.; Pfaender, S.; Ludwig, A. Nanoscale copper and silver thin film systems display differences in antiviral and antibacterial properties. Scientific Reports 2022, 12 (1), 10, Article. DOI: 10.1038/s41598-022- 11212-w.

(11) Godoy-Gallardo, M.; Eckhard, U.; Delgado, L. M.; de Roo Puente, Y. J.; Hoyos-Nogués, M.; Gil, F. J.; Perez, R. A. J. B. M. Antibacterial approaches in tissue engineering using metal ions and nanoparticles: From mechanisms to applications. 2021, 6 (12), 4470–4490.

(12) Merker, D.; Popova, B.; Bergfeldt, T.; Weingartner, T.; Braus, G. H.; Reithmaier, J. P.; Popov, C. Antimicrobial propensity of ultrananocrystalline diamond films with embedded silver nanodroplets. Diamond and Related Materials 2019, 93, 168–178, Article. DOI: 10.1016/j.diamond.2019.02.003.

(13) Tamulevicius, S.; Meskinis, S.; Tamulevicius, T.; Rubahn, H. G. Diamond like carbon nanocomposites with embedded metallic nanoparticles. Reports on Progress in Physics 2018, 81 (2), 31, Review. DOI: 10.1088/1361-6633/aa966f.

(14) Meskinis, S.; Ciegis, A.; Vasiliauskas, A.; Slapikas, K.; Tamulevicius, T.; Tamuleviciene, A.; Tamulevicius, S. Optical properties of diamond like carbon films containing copper, grown by high power pulsed magnetron sputtering and direct current magnetron sputtering: Structure and composition effects. Thin Solid Films 2015, 581, 48–53. DOI: 10.1016/j.tsf.2014.11.045.

(15) Khamseh, S.; Alibakhshi, E.; Mandavian, M.; Saeb, M. R.; Vahabi, H.; Kokanyan, N.; Laheurte, P. Magnetron-sputtered copper/diamond-like carbon composite thin films with super anti-corrosion properties. Surface & Coatings Technology 2018, 333, 148–157, Article. DOI: 10.1016/j.surfcoat.2017.11.012.

(16) Lan, W. C.; Ou, S. F.; Lin, M. H.; Ou, K. L.; Tsai, M. Y. Development of silver-containing diamond-like carbon for biomedical applications. Part I: Microstructure characteristics, mechanical properties and antibacterial mechanisms. Ceramics International 2013, 39 (4), 4099–4104, Article. DOI: 10.1016/j.ceramint.2012.10.264.

(17) Yan, C. L.; Guo, P.; Zhou, J. Y.; Chen, R. D.; Wang, A. Y. Dependence of piezoresistive behavior upon Cu content in Cu-DLC nanocomposite films. Diamond and Related Materials 2023, 136, 10, Article. DOI: 10.1016/j.diamond.2023.109935.

(18) Jurkevičiūtė, A.; Dolmantas, P.; Vasiliauskas, A.; Tamulevičienė, A.; Meškinis, Š.; Poplausks, R.; Prikulis, J.; Tamulevičius, S.; Tamulevičius, T. J. M. C.; Physics. Magnetron sputtering process for deposition of multilayered thin diamond-like carbon films with silver nanoparticles for anti-reflective coatings and refractometric sensing. 2023, 309, 128425.

(19) Thukkaram, M.; Vaidulych, M.; Kylián, O.; Rigole, P.; Aliakbarshirazi, S.; Asadian, M.; Nikiforov, A.; Biederman, H.; Coenye, T.; Du Laing, G. J. M. S.; et al. Biological activity and antimicrobial property of Cu/aC: H nanocomposites and nanolayered coatings on titanium substrates. 2021, 119, 111513.

(20) Zeng, Q.; Ning, Z. J. R. o. A. M. S. High-temperature tribological properties of diamond-like carbon films: A review. 2021, 60 (1), 276–292.

(21) Koutsokeras, L.; Constantinou, M.; Nikolaou, P.; Constantinides, G.; Kelires, P. J. M. Synthesis and characterization of hydrogenated diamond-like carbon (HDLC) nanocomposite films with metal (Ag, Cu) nanoparticles. 2020, 13 (7), 1753.

(22) Tamulevičius, S.; Meškinis, Š.; Tamulevičius, T.; Rubahn, H.-G. J. R. o. P. i. P. Diamond like carbon nanocomposites with embedded metallic nanoparticles. 2018, 81 (2), 024501.

(23) Chen, B.; Yan, F.; Yan, M.; Zhang, Y.; Xu, Y. J. A. A. M.; Interfaces. Experimental study of diamond-like carbon film on Aermet100 steel and first-principles calculation of interfacial adhesion. 2022, 14 (42), 48262–48275.

(24) Ji, L.; Li, H.; Zhao, F.; Chen, J.; Zhou, H. J. D.; Materials, R. Microstructure and mechanical properties of Mo/DLC nanocomposite films. 2008, 17 (11), 1949–1954.

(25) Javidparvar, A. A.; Mosavi, M. A.; Ramezanzadeh, B. J. M. C.; Physics. Nickel-aluminium bronze (NiBRAl) casting alloy tribological/corrosion resistance properties improvement via deposition of a Cu-doped diamond-like carbon (DLC) thin film; optimization of sputtering magnetron process conditions. 2023, 296, 127279.

(26) Liu, Y.; Guo, P.; He, X. Y.; Li, L.; Wang, A. Y.; Li, H. Developing transparent copper-doped diamond-like carbon films for marine antifouling applications. Diamond and Related Materials 2016, 69, 144–151, Article. DOI: 10.1016/j.diamond.2016.08.012.

(27) Juknius, T.; Ruzauskas, M.; Tamulevicius, T.; Siugzdiniene, R.; Jukniene, I.; Vasiliauskas, A.; Jurkeviciute, A.; Tamulevicius, S. Antimicrobial Properties of Diamond-Like Carbon/Silver Nanocomposite Thin Films Deposited on Textiles: Towards Smart Bandages. Materials 2016, 9 (5). DOI: 10.3390/ma9050371.

(28) Tamayo, L. A.; Zapata, P. A.; Vejar, N. D.; Azocar, M. I.; Gulppi, M. A.; Zhou, X.; Thompson, G. E.; Rabagliati, F. M.; Paez, M. A. Release of silver and copper nanoparticles from polyethylene nanocomposites and their penetration into Listeria monocytogenes. Materials Science and Engineering C-Materials for Biological Applications 2014, 40, 24–31, Article. DOI: 10.1016/j.msec.2014.03.037.

(29) Wu, X. H.; Ye, L.; Liu, K.; Wang, W.; Wei, J.; Chen, F. P.; Liu, C. S. Antibacterial properties of mesoporous copper-doped silica xerogels. Biomedical Materials 2009, 4 (4), 6, Article. DOI: 10.1088/1748-6041/4/4/045008.

(30) Joseph, L. A.; Israel, O. K.; Edet, E. J. Comparative Evaluation of Metal Ions Release from Titanium and Ti-6Al-7Nb into Bio-Fluids. 2009 2009, 6 (1).

(31) Mutlu-Sagesen, L.; Ergun, G.; Karabulut, E. Ion release from metal-ceramic alloys in three different media. Dental Materials Journal 2011, 30 (5), 598–610, Article. DOI: 10.4012/dmj.2011-031.

(32) Elguindi, J.; Moffitt, S.; Hasman, H.; Andrade, C.; Raghavan, S.; Rensing, C. J. A. m.; biotechnology. Metallic copper corrosion rates, moisture content, and growth medium influence survival of copper ion-resistant bacteria. 2011, 89, 1963–1970.

(33) Midander, K.; Cronholm, P.; Karlsson, H. L.; Elihn, K.; Möller, L.; Leygraf, C.; Wallinder, I. O. J. S. Surface characteristics, copper release, and toxicity of nano-and micrometer-sized copper and copper (II) oxide particles: a cross-disciplinary study. 2009, 5 (3), 389–399.

(34) Yang, K.; Chen, J.; Zheng, L.; Zheng, B.; Chen, Y.; Chen, X.; Bai, W.; Jian, R.; Wei, F.; Xu, Y. J. J. o. A. P. S. Urushiol titanium polymer-based composites coatings for anti-corrosion and antifouling in marine spray splash zones. 2021, 138 (34), 50861.

(35) Benhacine, F.; Hadj-Hamou, A. S.; Habi, A. Development of long-term antimicrobial poly (epsilon-caprolactone)/silver exchanged montmorillonite nanocomposite films with silver ion release property for active packaging use. Polymer Bulletin 2016, 73 (5), 1207–1227, Article. DOI: 10.1007/s00289-015-1543-9.

(36) Zakariene, G.; Novoslavskij, A.; Meskinis, S.; Vasiliauskas, A.; Tamuleviciene, A.; Tamulevicius, S.; Alter, T.; Malakauskas, M. Diamond like carbon Ag nanocomposites as a control measure against Campylobacter jejuni and Listeria monocytogenes on food preparation surfaces. Diamond and Related Materials 2018, 81, 118–126, Article. DOI: 10.1016/j.diamond.2017.12.007.

(37) Juknius, T.; Jukniene, I.; Tamulevicius, T.; Ruzauskas, M.; Pampariene, I.; Oberauskas, V.; Jurkeviciute, A.; Vasiliauskas, A.; Tamulevicius, S. Preclinical Study of a Multi-Layered Antimicrobial Patch Based on Thin Nanocomposite Amorphous Diamond Like Carbon Films with Embedded Silver Nanoparticles. Materials 2020, 13 (14), 14, Article. DOI: 10.3390/ma13143180.

(38) Tamulevicius, T.; Tamuleviciene, A.; Virganavicius, D.; Vasiliauskas, A.; Kopustinskas, V.; Meskinis, S.; Tamulevicius, S. Structuring of DLC:Ag nanocomposite thin films employing plasma chemical etching and ion sputtering. Nuclear Instruments & Methods in Physics Research Section B-Beam Interactions with Materials and Atoms 2014, 341, 1–6. DOI: 10.1016/j.nimb.2013.09.052.

(39) Purwar, T.; Esquivel-Puentes, H. A.; Pulletikurthi, V.; Li, X.; Doosttalab, A.; Nelson, C. E.; Appiah, R. E.; Blatchley, E. R.; Castano, V.; Castillo, L. Novel sustainable filter for virus filtration and inactivation. Scientific Reports 2022, 12 (1), 9, Article. DOI: 10.1038/s41598-022-13316-9.

(40) Meskinis, S.; Kopustinskas, V.; Tamuleviciene, A.; Tamulevicius, S.; Niaura, G.; Jankauskas, J.; Gudaitis, R. Ion beam energy effects on structure and properties of diamond like carbon films deposited by closed drift ion source. Vacuum 2010, 84 (9), 1133–1137, Article. DOI: 10.1016/j.vacuum.2010.01.047.

(41) Meškinis, Š.; Čiegis, A.; Vasiliauskas, A.; Šlapikas, K.; Tamulevičius, T.; Tamulevičienė, A.; Tamulevičius, S. J. T. S. F. Optical properties of diamond like carbon films containing copper, grown by high power pulsed magnetron sputtering and direct current magnetron sputtering: Structure and composition effects. 2015, 581, 48–53.

(42) Klinger, M. J. J. o. A. C. More features, more tools, more CrysTBox. 2017, 50 (4), 1226–1234.

(43) Mehanna, Y. A.; Crick, C. R. J. A. M. T. Image analysis methodology for a quantitative evaluation of coating abrasion resistance. 2021, 25, 101203.

44. Shanmugam, P. S. T.; Sampath, T.; Jagadeeswaran, I. Biocompatibility protocols for medical devices and materials; Elsevier, 2023.

(45) Lim, S.; Yap, A.; Loo, C.; Ng, J.; Goh, C.; Hong, C.; Toh, W. J. H.; toxicology, e. Comparison of cytotoxicity test models for evaluating resin-based composites. 2017, 36 (4), 339–348.

(46) Standardization, I. O. f. ISO 10993-5: 2009-Biological evaluation of medical devices-Part 5: Tests for in vitro cytotoxicity. 2009.

(47) Raffay, R.; Husin, N.; Omar, A. F. J. C. T. Spectrophotometry and colorimetry profiling of pure phenol red and cell culture medium on pH variation. 2022, 138 (6), 640–659.

(48) Mosmann, T. J. J. o. i. m. Rapid colorimetric assay for cellular growth and survival: application to proliferation and cytotoxicity assays. 1983, 65 (1-2), 55–63.

(49) Lisov, A.; Vrublevskaya, V.; Lisova, Z.; Leontievsky, A.; Morenkov, O. A 2,5- Dihydroxybenzoic Acid-Gelatin Conjugate: The Synthesis, Antiviral Activity and Mechanism of Antiviral Action Against Two Alphaherpesviruses. Viruses-Basel 2015, 7 (10), 5343–5360, Article. DOI: 10.3390/v7102878.

50. TCID50 calculator v2 17 01 20 MB - Excel sheet to calculate TCID50 titers (Spearman & Kärber method) by Marco Binder. https://www.klinikum.uni-heidelberg.de/zentrum-fuer-infektiologie/molecular-virology/welcome/downloads (accessed 2023 May 11, 2023).

(51) Wulff, N. H.; Tzatzaris, M.; Young, P. J. Monte Carlo simulation of the Spearman-Kaerber TCID50. Journal of Clinical Bioinformatics 2012, 2 (1), 5. DOI: 10.1186/2043-9113-2-5.

(52) Meir, R.; Maharat, O.; Farnushi, Y.; Simanov, L. Development of a real-time TaqMan (R) RT-PCR assay for the detection of infectious bronchitis virus in chickens, and comparison of RT-PCR and virus isolation. Journal of Virological Methods 2010, 163 (2), 190–194, Article. DOI: 10.1016/j.jviromet.2009.09.014.

(53) Abril, C.; Engels, M.; Liman, A.; Hilbe, M.; Albini, S.; Franchini, M.; Suter, M.; Ackermann, M. Both viral and host factors contribute to neurovirulence of bovine herpesviruses 1 and 5 in interferon receptor-deficient mice. Journal of Virology 2004, 78 (7), 3644–3653, Article. DOI: 10.1128/jvi.78.7.3644-3653.2004.

(54) Jurkeviciute, A.; Lazauskas, A.; Tamulevicius, T.; Vasiliauskas, A.; Peckus, D.; Meskinis, S.; Tamulevicius, S. Structure and density profile of diamond-like carbon films containing copper: Study by X-ray reflectivity, transmission electron microscopy, and spectroscopic ellipsometry. Thin Solid Films 2017, 630, 48–58, Article. DOI: 10.1016/j.tsf.2016.10.015.

(55) Kairaitis, G.; Galdikas, A. Mechanisms and Dynamics of Layered Structure Formation During Co-Deposition of Binary Compound Thin Films. Coatings 2020, 10 (1), 18, Article. DOI: 10.3390/coatings10010021.

56. Wagner, C. D. J. N. S. R. D. NIST X-ray photoelectron spectroscopy database. 2000.

(57) Biesinger, M. C.; Lau, L. W.; Gerson, A. R.; Smart, R. S. C. J. A. s. s. Resolving surface chemical states in XPS analysis of first row transition metals, oxides and hydroxides: Sc, Ti, V, Cu and Zn. 2010, 257 (3), 887–898.

58. 15472:, I. Surface Chemical Analysis—X-ray Photoelectron Spectrometers—Calibration of Energy Scales. ISO Geneva: 2010.

(59) Biesinger, M. C. J. S.; Analysis, I. Advanced analysis of copper X-ray photoelectron spectra. 2017, 49 (13), 1325–1334.

(60) Meškinis, Š.; Vasiliauskas, A.; Viskontas, K.; Andrulevičius, M.; Guobienė, A.; Tamulevičius, S. J. M. Hydrogen-free diamond like carbon films with embedded Cu nanoparticles: structure, composition and reverse saturable absorption effect. 2020, 13 (3), 760.

(61) Fredj, N.; Burleigh, T. D. Transpassive Dissolution of Copper and Rapid Formation of Brilliant Colored Copper Oxide Films. Journal of the Electrochemical Society 2011, 158 (4), C104–C110, Article. DOI: 10.1149/1.3551525.

(62) Nassau, K.; Gallagher, P. K.; Miller, A. E.; Graedel, T. E. The characterization of patina components by X-ray diffraction and evolved gas analysis. Corrosion Science 1987, 27 (7), 669–684. DOI: 10.1016/0010-938X(87)90049-7.

(63) Adeloju, s. B.; Duan, Y. Y. Corrosion resistance of CU2O and CuO on copper surfaces in aqueous media. British Corrosion Journal 1994, 29 (4), 309–314. DOI: 10.1179/000705994798267485.

(64) Nakayama, S.; Notoya, T.; Osakai, T. Highly Selective Determination of Copper Corrosion Products by Voltammetric Reduction in a Strongly Alkaline Electrolyte. Analytical Sciences 2012, 28 (4), 323–331, Review. DOI: 10.2116/analsci.28.323.

(65) Jiang, S.; D’Amario, L.; Dau, H. Copper Carbonate Hydroxide as Precursor of Interfacial CO in CO2 Electroreduction. Chemsuschem 2022, 15 (8), 13, Article. DOI: 10.1002/cssc.202102506.

(66) Yun, D. Y.; Choi, W. S.; Park, Y. S.; Hong, B. Effect of H-2 and O-2 plasma etching treatment on the surface of diamond-like carbon thin film. Applied Surface Science 2008, 254 (23), 7925–7928. DOI: 10.1016/j.apsusc.2008.03.170.

(67) Ferrari, A. C.; Robertson, J. Interpretation of Raman spectra of disordered and amorphous carbon. Physical Review B 2000, 61 (20), 14095–14107, Article. DOI: 10.1103/PhysRevB.61.14095.

(68) Ferrari, A. C.; Robertson, J. Raman spectroscopy of amorphous, nanostructured, diamond-like carbon, and nanodiamond. Philosophical Transactions of the Royal Society a-Mathematical Physical and Engineering Sciences 2004, 362 (1824), 2477–2512, Article. DOI: 10.1098/rsta.2004.1452.

(69) Dwivedi, N.; Kumar, S.; Malik, H. K.; Sreekumar, C.; Dayal, S.; Rauthan, C. M. S.; Panwar, O. S. Investigation of properties of Cu containing DLC films produced by PECVD process. Journal of Physics and Chemistry of Solids 2012, 73 (2), 308–316, Article. DOI: 10.1016/j.jpcs.2011.10.019.

(70) Dai, W.; Wang, A. Y.; Wang, Q. M. Microstructure and mechanical property of diamond-like carbon films with ductile copper incorporation. Surface & Coatings Technology 2015, 272, 33–38, Article. DOI: 10.1016/j.surfcoat.2015.04.027.

(71) Wei, X. J.; Yang, Z. D.; Wang, Y. X.; Tay, S. L.; Gao, W. Polymer antimicrobial coatings with embedded fine Cu and Cu salt particles. Applied Microbiology and Biotechnology 2014, 98 (14), 6265–6274, Article. DOI: 10.1007/s00253-014-5670-2.

(72) Quezada, R.; Quintero, Y.; Salgado, J. C.; Estay, H.; Garcia, A. Understanding the Phenomenon of Copper Ions Release from Copper-Modified TFC Membranes: A Mathematical and Experimental Methodology Using Shrinking Core Model. Nanomaterials 2020, 10 (6), 18, Article. DOI: 10.3390/nano10061130.

(73) Chatterjee, A. K.; Chakraborty, R.; Basu, T. Mechanism of antibacterial activity of copper nanoparticles. Nanotechnology 2014, 25 (13), 12, Article. DOI: 10.1088/0957-4484/25/13/135101.

(74) Warnes, S. L.; Summersgill, E. N.; Keevil, C. W. Inactivation of Murine Norovirus on a Range of Copper Alloy Surfaces Is Accompanied by Loss of Capsid Integrity. Applied and Environmental Microbiology 2015, 81 (3), 1085–1091, Article. DOI: 10.1128/aem.03280-14.

(75) Merkl, P.; Long, S. W.; McInerney, G. M.; Sotiriou, G. A. Antiviral Activity of Silver, Copper Oxide and Zinc Oxide Nanoparticle Coatings against SARS-CoV-2. Nanomaterials 2021, 11 (5), 9, Article. DOI: 10.3390/nano11051312.

(76) Jablonská, E.; Kubásek, J.; Vojtěch, D.; Ruml, T.; Lipov, J. J. S. R. Test conditions can significantly affect the results of in vitro cytotoxicity testing of degradable metallic biomaterials. 2021, 11 (1), 6628.

(77) Shigetoh, K.; Hirao, R.; Ishida, N. J. A. a. m.; interfaces. Durability and surface oxidation states of antiviral nano-columnar copper thin films. 2023, 15 (16), 20398–20409.

(78) Sagripanti, J. L.; Routson, L. B.; Bonifacino, A. C.; Lytle, C. D. Mechanism of copper-mediated inactivation of herpes simplex virus. Antimicrobial Agents and Chemotherapy 1997, 41 (4), 812–817, Article. DOI: 10.1128/aac.41.4.812.

(79) Ishida, T. Antiviral Activities of Cu2+ Ions in Viral Prevention, Replication, RNA Degradation, and for Antiviral Efficacies of Lytic Virus, ROS-Mediated Virus, Copper Chelation. World Scientific News 2018, 99, 148–168.

(80) Mantlo, E. K.; Paessler, S.; Seregin, A.; Mitchell, A. Luminore CopperTouch Surface Coating Effectively Inactivates SARS-CoV-2, Ebola Virus, and Marburg Virus In Vitro. Antimicrobial Agents and Chemotherapy 2021, 65 (7), 5, Article. DOI: 10.1128/aac.01390-20.

(81) Behzadinasab, S.; Chin, A.; Hosseini, M.; Poon, L.; Ducker, W. A. A Surface Coating that Rapidly Inactivates SARS-CoV-2. Acs Applied Materials & Interfaces 2020, 12 (31), 34723–34727, Article. DOI: 10.1021/acsami.0c11425.

(82) Gurunathan, S.; Qasim, M.; Choi, Y.; Do, J. T.; Park, C.; Hong, K.; Kim, J. H.; Song, H. Antiviral Potential of Nanoparticles-Can Nanoparticles Fight Against Coronaviruses? Nanomaterials 2020, 10 (9), 29, Review. DOI: 10.3390/nano10091645.

(83) Hutasoit, N.; Kennedy, B.; Hamilton, S.; Luttick, A.; Rashid, R. A. R.; Palanisamy, S. Sars-CoV-2 (COVID-19) inactivation capability of copper-coated touch surface fabricated by cold-spray technology. Manufacturing Letters 2020, 25, 93–97, Article. DOI: 10.1016/j.mfglet.2020.08.007.

(84) Thevenon, F.; Regier, N.; Benagli, C.; Tonolla, M.; Adatte, T.; Wildi, W.; Pote, J. Characterization of fecal indicator bacteria in sediments cores from the largest freshwater lake of Western Europe (Lake Geneva, Switzerland). Ecotoxicology and Environmental Safety 2012, 78, 50–56, Article. DOI: 10.1016/j.ecoenv.2011.11.005.

(85) Li, X. M.; Cong, Y. L.; Ovais, M.; Cardoso, M. B.; Hameed, S.; Chen, R.; Chen, M. L.; Wang, L. M. Copper-based nanoparticles against microbial infections. Wiley Interdisciplinary Reviews- Nanomedicine and Nanobiotechnology 2023, 15 (4), 25, Review. DOI: 10.1002/wnan.1888.

(86) Liu, M.; Gong, X.; Alluri, R. K.; Wu, J. H.; Sablo, T.; Li, Z. W. Characterization of RNA damage under oxidative stress in Escherichia coli. Biological Chemistry 2012, 393 (3), 123–132, Article. DOI: 10.1515/hsz-2011-0247.

