## Supplementary figures and images for "Antiviral and Antibacterial Efficacy of Nanocomposite Amorphous Carbon Films with Copper Nanoparticles"

### Graphical abstract

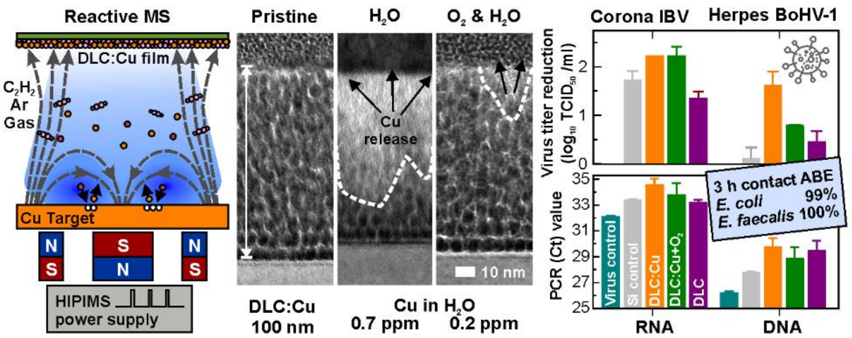
