## Supplementary information for "Antiviral and Antibacterial Efficacy of Nanocomposite Amorphous Carbon Films with Copper Nanoparticles"

The half-life for the viruses and bacteria was calculated based on **eq. S1** [1]:

$$N(t) = N_0 \left(\frac{1}{2}\right)^{\frac{t}{t_{1/2}}} \implies t_{1/2} = \frac{-ln(2)\cdot t}{ln\left(\frac{N(t)}{N_0}\right)} \qquad \textbf{(eq. S1)}$$

where $N(t)$ – remaining quantity, $N_0$ – initial quantity, $t$ – elapsed time, $t_{1/2}$ – half-life of the substance.

The antibacterial effectiveness (ABE, %) was calculated using **eq. S2**:

$$ABE(\%) = \frac{(N_{ref} - N_{exp})}{N_{ref}} \cdot 100\% \qquad \textbf{(eq. S2)}$$

where $N_{ref}$ - is the number of bacteria in the control plate sample, and $N_{exp}$ - is the number of bacteria in the sample of the experimental plate.

The original methodology developed for the inspection of the DLC:Cu film virucidal efficacy degradation after its immersion in the cell medium is depicted in **Table S1**

**Table S1** Methodology for DLC:Cu film virucidal efficacy stability evaluation over prolonged time periods.

| **Incubation duration and testing** | **Medium type** | | | | | | |
|---|---|---|---|---|---|---|---|
| | Virus | Cell medium | Virus | Cell medium | Virus | Cell medium | Virus |
| Duration of individual treatment (h) | 1 | 2 | 1 | 4 | 1 | 15 | 1 |
| Overall accumulated exposure duration (h) | 1 | 3 | 4 | 8 | 9 | 24 | 25 |

The energy dispersive x-ray spectroscopy determined elemental analysis results of pristine and plasma-processed as well as ultra-pure water-immersed DLC:Cu films are tabulated in **Table S2**.

**Table S2** Pristine DLC:Cu and different duration $O_2$ plasma processed film average elemental composition before and after exposure to ultra-pure water. The error bars stand for one standard deviation.

| **Elements** | **Concentration (at.%)** | | | | | |
|---|---|---|---|---|---|---|
| **Duration of $O_2$ plasma processing (s)** | - | 5 | 20 | - | 5 | 20 |
| **Immersion in water (h/°C)** | - | - | - | 24/37 | 24/37 | 24/37 |
| **Carbon (C)** | 34.8±0.9 | 34.2±0.6 | 35.2±2.5 | 41.3±2.0 | 22.3±1.0 | 24.6±0.5 |
| **Oxygen (O)** | 3.8±0.4 | 7.0±0.9 | 5.7±2.2 | 6.6±1.7 | 37.1±1.9 | 30.5±0.1 |
| **Copper (Cu)** | 61.4±0.7 | 58.8±0.3 | 59.1±0.5 | 52.0±0.7 | 40.6±1.0 | 44.9±0.4 |

Comparison of the XPS spectra in the Cu 2p region for all samples is presented in **Figure S1 a**. All samples show obvious similarity of the spectra in this region except the sample exposed to oxygen plasma for 5 s and immersed in water for 24 hours (**Figure S1 a**). For this sample appearance of the clear satellite peaks at 940 eV and 945 eV [2] indicates that Cu(II) is present in $Cu(OH)_2$ or/and CuO bonds.

X-rays excited Cu LMM spectra for the same samples are shown in **Figure S1 b**. In this picture, the absence of a metallic copper peak at 568 eV indicates that copper on the surface of the samples is in the oxidized state. For most samples, the main peak is located approximately at 659.9 eV, corresponding to the known position (569.89 eV, 569.61 eV) [2, 3] of $Cu_2O$ bonds. The main peak (at 569.2 eV) of the sample exposed to oxygen plasma for 5 s and immersed in water for 24 hours (**Figure S1 b**) is clearly shifted towards the CuO position (568.7 eV). This supports the presence of Cu(II) in CuO bonds as was shown in the Cu 2p region also, in the text above. The sample treated in $O_2$ plasma for 5 s (**Figure S1 b**) shows a narrow peak at 569.9 eV typical for $Cu_2O$ bonds. The sample immersed in virus solution for 24 hours (**Figure S1 b**) also shows the main peak at 569.9 eV (typical for $Cu_2O$ bonds) but the peak is wider and indicates the presence of $Cu(OH)_2$ bonds (570.4 eV, 570.35 eV) [2, 3] in the sample.

The XPS spectra in the O 1s region for the same samples are presented in **Figure S1 c**. In this graph the spectra for plasma treated and immersed in pure water sample show the presence of Cu(II) in CuO (529.8 eV), Cu(I) in $Cu_2O$ (530.5 eV) [4], and some amount of C=O (531.6 eV), C-O (532.8 eV) [3] bonds as well. Similar to the findings in the Cu LMM region, all other samples demonstrated $Cu_2O$ (530.5 eV) bonds and C=O, C-O bonds. The presence of $Cu(OH)_2$ bonds can be excluded for all samples, as was shown in the Cu LMM region, except for the sample immersed in virus solution.

The XPS spectra in the C 1s region for the same samples are presented in **Figure S1 d**. In this graph, almost identical carbon C 1s spectra (**Figure S1 d**) for all samples are demonstrated, showing that most carbon atoms are present in C-C bonds ($sp^3$ hybridization), typical for DLC films. The small amount of carbon in C=C bonds ($sp^2$) and carbon-oxygen (C-O, C=O, O=C-O) bonds is also present, as was found in the O 1s region.

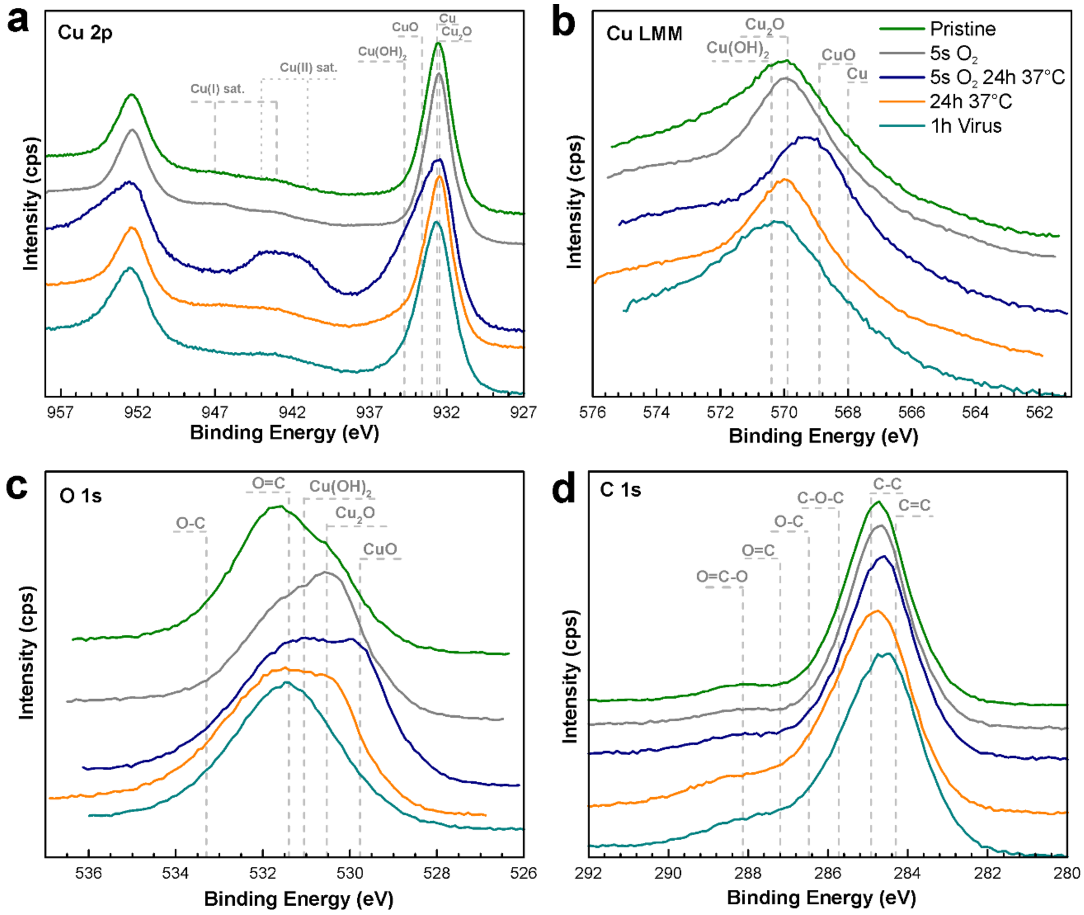


**Figure S1** XPS spectra comparison for DLC:Cu samples before and after treatment in plasma and immersing in water in different binding energy regions. (a) Cu 2p region. (b) Copper Auger spectra in Cu LMM region. (c) O 1s region. (d) C 1s region.

The Cu compound release in ultra-pure water of pristine and plasma-preprocessed DLC:Cu films obtained with atomic absorption spectroscopy are tabulated in **Table S3**.

**Table S3** AAS measured Cu concentration normalized to 5 ml solvent and 1 $cm^2$ area released from pristine and different duration plasma processed DLC:Cu films in ultra-pure water incubated at different temperatures and durations.

| **Temperature (°C)** | **Released Cu concentration (mg/L/$cm^2$ or ppm/$cm^2$)** | | | | | |
|---|---|---|---|---|---|---|
| **$O_2$ processing duration (s)** | - | | 5 | | 20 | |
| **Exposure to water duration (h)** | 24 | 168 | 24 | 168 | 24 | 168 |
| **5** | 0.047* | 1.333 | 0.094* | 1.009 | 0.136* | 0.967 |
| **37** | 0.507 | 0.737 | 0.277 | 0.277 | 0.183 | 0.324 |

*Results close to the limit of detection.

The optical microscope images of the DLC:Cu films after abrasion with different sandpaper grits, rotations per minute, and rotation cycles are summarized in **Figure S2 a-b** while the impacted area analysis is depicted in **Figure S2 g**. The area analysis was carried out employing ImageJ software.

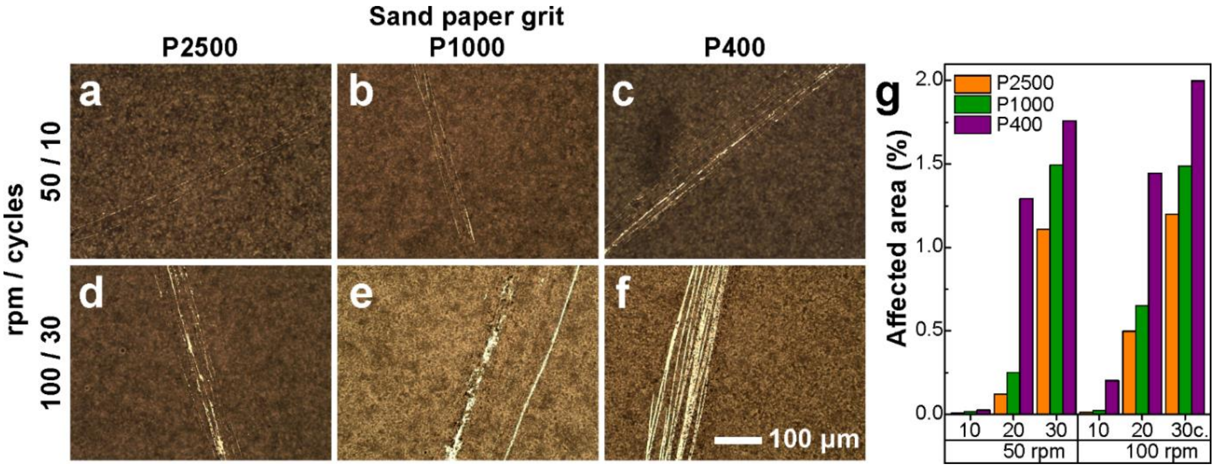


**Figure S2** Optical microscope images of the abraded area on the surface of the DLC:Cu film after each abrasion cycle with different sandpapers grit (a,b,c) 50 rpm/ 10 cycle and (d,e,f) 100 rpm/ 30 cycle, (g) a plot of the percentage of the affected area.

The colours of the DLC:Cu films visible by the naked eye after exposure to ultra-pure water are depicted in the sample images in **Figure S3**.

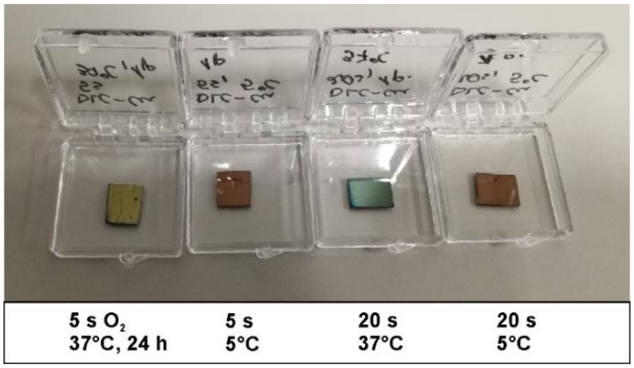


**Figure S3** Camera images of 5 s and 20 s plasma processed DLC:Cu samples after immersion for 24 hours in 5ºC and 37ºC temperatures as indicated below the samples. Sample edge length *ca.* 1 cm.

For a better impression of the Cu release from the DLC:Cu in the virucidal study the preliminary analysis of the aqueous medium volume and thickness above the immersed sample influence on the released Cu concentration was conducted in shorter immersion durations (5 min, 10 min, 15 min, 60 min) using Dulbecco's modified Eagle's medium (DMEM, Gibco, UK). The same volume of 5 ml was used and two different film area samples, namely 2.1 cm$^2$ and 55.1 cm$^2$, were immersed in different size Petri dishes resulting in different medium volumes and effective thicknesses above the sample.

The same double-beam atomic absorption spectrometer (AAS) AAnalyst 400 (Perkin Elmer, USA) was used as explained in the main text. The AAS measurement results depicted in **Figure S3** indicate that accumulated Cu concentration in the medium is bigger and approaching 2.5 mg/L/cm$^2$ when the liquid volume was relatively big compared to the sample area. When the big area samples were used and the 5 ml aqueous medium was just covering the sample surface the Cu release was almost 10 times smaller in the same solvent volume than previously. The Cu release in ultra-pure water (**Figure 3 b, Table S3**) was slower than in DMEM **Figure S4**.

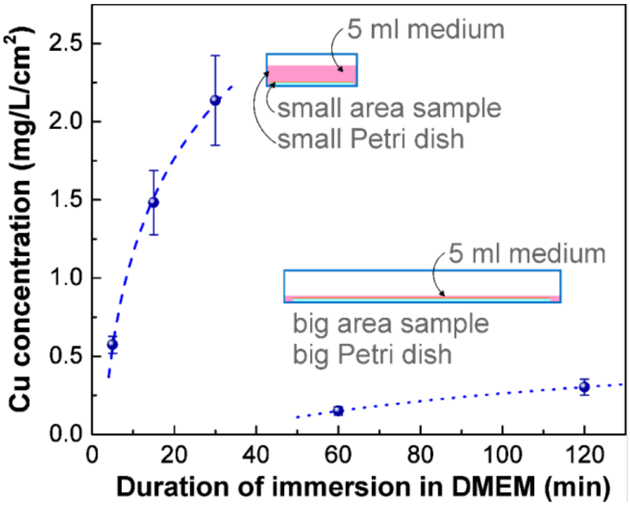


**Figure S4** Accumulated Cu release from pristine DLC:Cu films in 5 ml DMEM at room temperature after different exposure durations using 2.1 cm$^2$ and 55.1 cm$^2$ area film samples.

Cytotoxicity results of MDBK (**Table S4**) and Vero cells (**Table S5**) after incubation for 24 h in different dilutions of extract obtained after DLC:Cu immersion in cell culture media with 5% FBS at 37°C for 24 hours. The estimated $CC_{50}$ for MDBK cells was calculated at a dilution of 1:9.71 (3.28 $log_2$). The estimated $CC_{50}$ for Vero cells was calculated at a dilution of 1:3.09 (1.75 $log_2$).

**Table S4** The cytotoxicity of extract of DLC:Cu for MDBK cells after the incubation of 24 hours

| Cell culture and extract concentration | Cell morphology, grade | Cells Viab, %±SD | pH |
|---|---|---|---|
| **MDBK/Cu 1:1 (100%)** | 4 | 7.7±0.73 | 7.7 |
| **MDBK/Cu 1:2 (50%)** | 4 | 13.8±1.12 | 7.7 |
| **MDBK/Cu 1:4 (25%)** | 3 | 19.4±1.69 | 7.6 |
| **MDBK/Cu 1:8 (12.5%)** | 2 | 46.3±4.98 | 7.3 |
| **MDBK/Cu 1:16 (6.3%)** | 1 | 83.2±4.96 | 7.3 |
| **MDBK/Cu 1:32 (3.2%)** | 1 | 103.5±4.72 | 7.3 |
| **MDBK/Cu 1:64 (1.6%)** | 0 | 101.0±4.27 | 7.3 |
| **MDBK/DLC 1:1 to 1:64 (100 to 1.6%)** | 0 | 100 | 7.3 |
| **MDBK negative control (blank)** | 0 | 99.9±3.64 | 7.3 |
| **MDBK positive control (Triton x 100)** | 4 | 1.6±0.29 | 7.4 |

**Table S5** The cytotoxicity of extract of DLC:Cu for Vero cells after the incubation of 24 hours

| Cell culture and extract concentration | Cell morphology, Grade | Cells Viab, %±SD | pH |
|---|---|---|---|
| **Vero/Cu 1:1 (100%)** | 4 | 14.7±3.00 | 7.7 |
| **Vero /Cu 1:2 (50%)** | 3 | 30.7±1.39 | 7.7 |
| **Vero K/Cu 1:4 (25%)** | 2 | 69.5±4.13 | 7.5 |
| **Vero K/Cu 1:8 (12.5%)** | 1 | 87.1±4.89 | 7.3 |
| **Vero /Cu 1:16 (6.3%)** | 1 | 100.4±3.97 | 7.3 |
| **Vero /Cu 1:32 (3.2%)** | 0 | 102.1±3.32 | 7.3 |
| **Vero /Cu 1:64 (1.6%)** | 0 | 100.7±3.10 | 7.3 |
| **Vero /DLC 1:1 to 1:64 (100 to 1.6%)** | 0 | 100 | 7.3 |
| **Vero negative control (blank)** | 0 | 100 | 7.3 |
| **Vero positive control (Triton x100)** | 4 | 2.2±0.33 | 7.4 |

Morphological changes of MDBK and VERO cells after being in contact with different dilution extracts obtained from DLC:Cu samples are depicted in **Figure S4.**

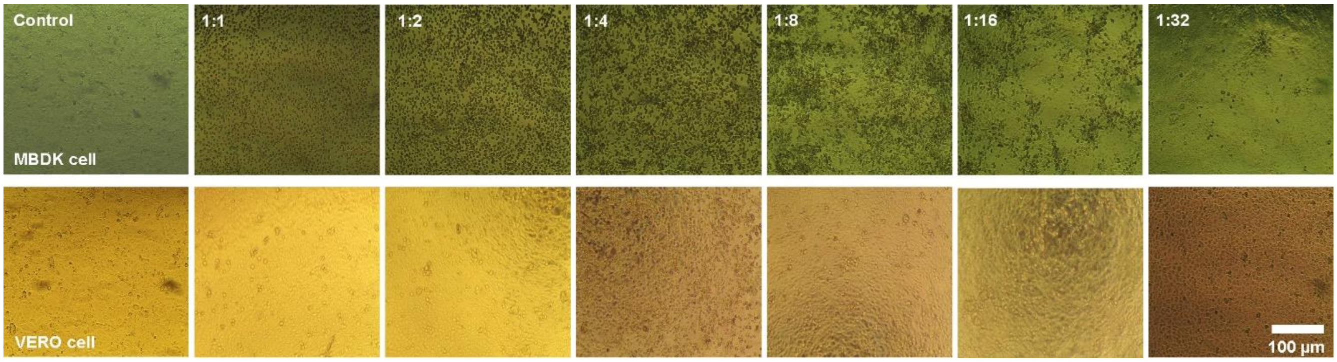

**Figure S4** Optical microscope images of the cytotoxicity test on the MDBK (top row) and VERO cell cultures (bottom row) showing the influence of DLC:Cu films on the cell morphology with different dilution ratios (indicated on the top left corner) with the cell culture medium.

Virucidal testing results of IBV and BoHV-1 are summarized in **Table S6** and **Table S7** respectively.

**Table S6** IBV strain ”Beaudette” activity after 1-hour contact with DLC:Cu, DLC:Cu+$O_2$, DLC films, and controls. 10 µl of the virus solution was used, and the temperature was 20±2°C. Washed for 1 min with 0.49 ml DMEM (initial dilution – 1:50).

| **Sample** | **Viral titre, $log_{10}$ $TCID_{50}$/ml, *M±σ*** | **Residual viruses, $TCID_{50}$ units/ml** | **A decrease in the number of viruses** | | **Half-life (min)** | **PCR $C_t$ value, *M±σ*** |
|---|---|---|---|---|---|---|
| | | | **$log_{10}$** | **%** | | |
| **IBV control** | 5.13±0.18 | 133 352 | - | - | - | 32.1±0.09 |
| **Si control** | 3.40±0.19 | 2 511 | 1.77 | 98.14 | 10.44 | 33.4±0.08 |
| **DLC:Cu** | 2.90±0.00 | 791 | 2.23 | 99.41 | 8.10 | 34.6±0.45 |
| **DLC:Cu+$O_2$** | 2.90±0.18 | 791 | 2.23 | 99.41 | 8.10 | 33.8±0.88 |
| **DLC** | 3.77±0.13 | 5 930 | 1.36 | 95.63 | 13.28 | 33.2±021 |

**Table S7** BoHV-1 strain ”4016” activity after 1-hour contact with DLC:Cu, DLC:Cu+$O_2$, DLC films, and controls. 10 µl of the virus solution was used, and the temperature was 20±2°C. Washed for 1 min with 0.49 ml DMEM/F12 (initial dilution – 1:50).

| **Sample** | **Viral titre, $log_{10}$ $TCID_{50}$/ml, *M±σ*** | **Residual viruses, $TCID_{50}$ units/ml** | **A decrease in the number of viruses** | | **Half-life (min)** | **PCR $C_t$ value, *M±σ*** |
|---|---|---|---|---|---|---|
| | | | **$log_{10}$ $TCID_{50}$/ml** | **%** | | |
| **BoHV-1 control** | 7.00±0.22 | 10 000 000 | - | - | - | 26.2±0.15 |
| **Si control** | 6.87±0.21 | 7 413 102 | 0.13 | 25.90 | 138.94 | 27.8±0.08 |
| **DLC:Cu** | 5.37±0.27 | 234 422 | 1.63 | 97.66 | 11.08 | 29.8±0.65 |
| **DLC:Cu+$O_2$** | 6.20±0.01 | 1 584 893 | 0.80 | 84.15 | 22.58 | 28.9±0.86 |
| **DLC*** | 6.53±0.21 | 3 388 441 | 0.64 | 66.11 | 38.43 | 29.5±0.75 |

*For DLC testing different control sample was used: 7.17±0.28 viral titre ($log_{10}$ $TCID_{50}$/ml).

Virucidal activity results of the DLC:Cu films inspected with BoHV – 1 and IBV are depicted in **Table S8** and **Table S9** respectively

**Table S8**. DLC:Cu surface activity dynamics and residual BoHV-1 titre $log_{10}$ after contact with Cu containing film.

| **The cycle of virus treatment** | **Hours of experiment** | **Titre $log_{10}$, *M±σ*** | **Decrease of titre, $log_{10}$** | **Decrease in number of viruses, %** |
|---|---|---|---|---|
| **1** | 0-1 | 4.25±0.25 | 3.00 | 99.90 |
| **2** | 3-4 | 4.95±0.25 | 2.30 | 99.50 |
| **3** | 8-9 | 5.45±0.25 | 1.80 | 98.42 |

| | | | | |
|---|---|---|---|---|
| **4** | 24-25 | 6.2±0 | 1.05 | 91.09 |
| **Control** | | 7.25±0.25 | - | - |

**Table S9**. DLC:Cu surface activity dynamics and residual IBV titre $log_{10}$ after contact with Cu containing film.

| **The cycle of virus treatment** | **Hours of experiment** | **Titre $log_{10}$, $M±\sigma$** | **Decrease of titre, $log_{10}$** | **Decrease in number of viruses, %** |
|---|---|---|---|---|
| **1** | 0-1 | 3.00±0.19 | 2.00 | 99.00 |
| **2** | 3-4 | 3.75±0.16 | 1.25 | 94.38 |
| **3** | 8-9 | 4.25±0.16 | 0.75 | 82.22 |
| **4** | 24-25 | 4.25±0.16 | 0.75 | 82.22 |
| **Control** | | 5.00±0.19 | - | - |

The antibacterial testing results that were repeated three times for each investigated *E. coli* and *E. faecalis* bacteria strains are tabulated in **Tables S10-S12** and **Tables S13-S15**, respectively.

**Table S10** Resulting number of colony forming units (CFU), and antibacterial efficiency against *E. coli* bacteria strain ATCC 25922 after being in contact with pristine DLC:Cu film and a silicon substrate. Test 1. Initial bacteria concentration 6.04 $log_{10}$ CFU/ml.

| **Samples** | | ***E. coli* number in PBS broth** | | | **Antibacterial efficiency (%)** | **Half-life (min)** |
|---|---|---|---|---|---|---|
| **Exposure duration (hours)** | **No.** | **$10^6$ CFU/ml** | **$10^6$ CFU/ml** | **$10^4$ CFU/ml** | | |
| | | **Si control** | **DLC** | **DLC:Cu** | | |
| **1** | **1** | 5.9 | 5.9 | 3.3 | 99.7 | 6.7 |
| | **2** | 5.9 | 5.9 | 3.3 | 99.7 | 6.7 |
| | **3** | 6 | 6 | 3.3 | 99.8 | 4.8 |
| Average: | | 5.98 | 5.98 | 3.3 | 99.79 | 6.10 |
| **3** | **1** | 5.9 | 5.8 | 0 | 100 | - |
| | **2** | 5.8 | 5.8 | 0 | 100 | - |
| | **3** | 5.8 | 5.8 | 0 | 100 | - |
| Average: | | 5.88 | 5.85 | 0 | 100.00±0.00 | - |
| **8** | **1** | 5.6 | 5.6 | 0 | 100 | - |
| | **2** | 5.96 | 5.6 | 0 | 100 | - |
| | **3** | 5.7 | 5.6 | 0 | 100 | - |
| Average: | | 5.72 | 5.66 | 0 | 100.00±0.00 | - |
| **24** | **1** | 5.8 | 6.6 | 0 | 100 | - |
| | **2** | 5.8 | 6.6 | 0 | 100 | - |
| | **3** | 5.8 | 6.6 | 0 | 100 | - |
| Average: | | 5.81 | 6.66 | 0 | 100.00±0.00 | - |

**Table S11**. The resulting number of CFU and antibacterial efficiency *E. coli* bacteria strain ATCC 25922 after being in contact with pristine DLC:Cu film and silicon substrate. Test 2. Initial bacteria concentration 6.0 $\log_{10}$ CFU/ml.

| Samples / Time (h) | No. | ***E. coli* number in PBS broth** $10^6$ CFU/ml Si control | $10^6$ CFU/ml DLC | $10^4$ CFU/ml DLC:Cu | Antibacterial efficiency (%) | Half-life (min) |
|---|---|---|---|---|---|---|
| **1** | 1 | 6 | 5.9 | 2 | 99.9 | 4.5 |
| | 2 | 5.9 | 5.9 | 2.1 | 99.9 | 4.6 |
| | 3 | 5.9 | 6 | 2 | 99.9 | 3.6 |
| Average: | | 5.98 | 5.97 | 2.02 | 99.98 | 4.25 |
| **3** | **1** | 5.9 | 5.9 | 0 | 100 | - |
| | **2** | 5.8 | 5.9 | 0 | 100 | - |
| | **3** | 5.9 | 5.9 | 0 | 100 | - |
| Average: | | 5.90 | 5.91 | 0 | 100.00±0.00 | - |
| **8** | **1** | 5.7 | 5.7 | 0 | 100 | - |
| | **2** | 5.67 | 5.7 | 0 | 100 | - |
| | **3** | 5.7 | 5.7 | 0 | 100 | - |
| Average: | | 5.70 | 5.75 | 0 | 100.00±0.00 | - |
| **24** | **1** | 5.7 | 6.6 | 0 | 100 | - |
| | **2** | 5.7 | 6.7 | 0 | 100 | - |
| | **3** | 5.6 | 6.7 | 0 | 100 | - |
| Average: | | 5.71 | 6.69 | 0 | 100.00±0.00 | - |

**Table S12** The resulting number of CFU, antibacterial efficiency, and half-time against *E. coli* strain ATCC 29212 bacteria after being in contact with pristine DLC:Cu film and silicon substrate. Test 3. Initial bacteria concentration 6.05 $\log_{10}$ CFU/ml.

| Samples / Time (h) | No. | ***E. coli* number in PBS broth** $10^6$ CFU/ml Si control | $10^6$ CFU/ml DLC | $10^4$ CFU/ml DLC:Cu | Antibacterial efficiency (%) | Half-life (min) |
|---|---|---|---|---|---|---|
| **1** | 1 | 5.9 | 5.9 | 0 | 100 | - |
| | 2 | 5.9 | 5.9 | 0 | 100 | - |
| | 3 | 5.9 | 6 | 0 | 100 | - |
| **Average:** | | 5.96 | 5.97 | 0 | 100.00±0.00 | - |
| **3** | **1** | 5.7 | 5.6 | 0 | 100 | - |
| | **2** | 5.7 | 5.7 | 0 | 100 | - |
| | **3** | 5.7 | 5.6 | 0 | 100 | - |
| | Average: | 5.71 | 5.69 | 0 | 100.00±0.00 | - |
| **8** | **1** | 5.5 | 5.4 | 0 | 100 | - |

| | | | | | | |
|---|---|---|---|---|---|---|
| | **2** | 5.5 | 5.4 | 0 | 100 | - |
| | **3** | 5.5 | 5.4 | 0 | 100 | - |
| Average: | | 5.57 | 5.45 | 0 | 100.00±0.00 | - |
| **24** | **1** | 5.5 | 5.4 | 0 | 100 | - |
| | **2** | 5.5 | 5.4 | 0 | 100 | - |
| | **3** | 5.5 | 5.4 | 0 | 100 | - |
| Average: | | 5.55 | 5.48 | 0 | 100.00±0.00 | - |

**Table S13** The resulting number of CFU, antibacterial efficiency, and half-time against *E. faecalis* strain ATCC 29212 bacteria after being in contact with pristine DLC:Cu film and silicon substrate. Test 1. Initial bacteria concentration 6.02 $\log_{10}$ CFU/ml.

| **Samples** | | ***E. faecalis* number in PBS broth** | | | **Antibacterial efficiency (%)** | **Half-life (min)** |
|---|---|---|---|---|---|---|
| **Exposure duration (hours)** | **No.** | **$10^6$ CFU/ml** | **$10^6$ CFU/ml** | **$10^4$ CFU/ml** | | |
| | | **Si control** | **DLC** | **DLC:Cu** | | |
| **1** | 1 | 5.8 | 5.8 | 5.7 | 15.06 | 254.6 |
| | 2 | 5.8 | 5.8 | 5.7 | 13.88 | 278.1 |
| | 3 | 5.8 | 5.8 | 5.8 | 6.66 | 602.7 |
| Average: | | 5.82 | 5.86 | 5.80 | 11.81 | 378.5 |
| **3** | **1** | 5.6 | 5.8 | 4.9 | 88.57 | 58 |
| | **2** | 5.6 | 5.7 | 4.9 | 85 | 65.7 |
| | **3** | 5.6 | 5.7 | 4.8 | 86.53 | 62.2 |
| Average: | | 5.60 | 5.76 | 4.88 | 86.87±1.7 | 61.9±3.8 |
| **8** | **1** | 5.6 | 5.6 | 0 | 100 | - |
| | **2** | 5.6 | 5.6 | 0 | 100 | - |
| | **3** | 5.6 | 5.6 | 0 | 100 | - |
| Average: | | 5.62 | 5.60 | 0 | 100.00±0.00 | - |
| **24** | **1** | 4.7 | 4.8 | 0 | 100 | - |
| | **2** | 4.7 | 4.8 | 0 | 100 | - |
| | **3** | 4.7 | 4.8 | 0 | 100 | - |
| | Average: | 4.76 | 4.81 | 0 | 100.00±0.00 | - |

**Table S14** Resulting number of CFU, antibacterial efficiency, and half-time against *E. faecalis* bacteria ATCC 29212 strain after being in contact with pristine DLC:Cu film and silicon substrate. Test 2. Initial bacteria concentration 6.04 $\log_{10}$ CFU/ml.

| **Samples** | | ***E. faecalis* number in PBS broth** | | | **Antibacterial efficiency (%)** | **Half-life (min)** |
|---|---|---|---|---|---|---|
| **Exposure duration (hours)** | **No.** | **$10^6$ CFU/ml** | **$10^6$ CFU/ml** | **$10^5$ CFU/ml** | | |
| | | **Si control** | **DLC** | **DLC:Cu** | | |
| **1** | **1** | 5.9 | 5.8 | 4.8 | 90.1 | 17.9 |
| | **2** | 5.9 | 5.9 | 4.8 | 92.3 | 16.9 |
| | **3** | 5.9 | 5.8 | 4.8 | 91.2 | 17.5 |
| Average | | 5.92 | 5.87 | 4.84 | 91.70 | 17.4 |

| | | | | | | |
|---|---|---|---|---|---|---|
| **3** | **1** | 5.8 | 5.8 | 2.2 | 99.9 | 15.06 |
| | **2** | 5.8 | 5.8 | 2.3 | 99.9 | 15.4 |
| | **3** | 5.8 | 5.8 | 2.2 | 99.9 | 15.1 |
| Average: | | 5.85 | 5.82 | 2.26 | 99.97±0.01 | 15.21±0.18 |
| **8** | **1** | 5.5 | 5.6 | 0 | 100 | |
| | **2** | 5.5 | 5.6 | 0 | 100 | |
| | **3** | 5.5 | 5.6 | 0 | 100 | |
| Average: | | 5.55 | 5.61 | 0 | 100.00±0.00 | |
| **24** | **1** | 4.98 | 5 | 0 | 100 | |
| | **2** | 5 | 5.1 | 0 | 100 | |
| | **3** | 4.9 | 5 | 0 | 100 | |
| Average: | | 4.98 | 5.06 | 0 | 100.00±0.00 | |

**Table S15** Resulting number of CFU, antibacterial efficiency, and half-time against *E. faecalis* bacteria ATCC 29212 strain after being in contact with pristine DLC:Cu film and silicon substrate. Test 3. Initial bacteria concentration 6.05 $log_{10}$ CFU/ml.

| **Samples** | | ***E. faecalis* number in PBS broth** | | | | |
|---|---|---|---|---|---|---|
| **Exposure duration (hours)** | **No.** | **$10^6$ CFU/ml** | **$10^6$ CFU/ml** | **$10^5$ CFU/ml** | **Antibacterial efficiency (%)** | **Half-life (min)** |
| | | **Si control** | **DLC** | **DLC:Cu** | | |
| **1** | **1** | 5.8 | 5.8 | 5.6 | 41.7 | 95.8 |
| | **2** | 5.8 | 5.9 | 5.6 | 38.3 | 88.4 |
| | **3** | 5.8 | 5.8 | 5.6 | 42.6 | 87.9 |
| Average | | 5.87 | 5.84 | 5.64 | 40.96 | 92.1 |
| **3** | **1** | 5.7 | 5.6 | 4.7 | 89.03 | 59.1 |
| | **2** | 5.7 | 5.6 | 4.7 | 89.8 | 62.8 |
| | **3** | 5.7 | 5.6 | 4.7 | 89.09 | 60 |
| Average: | | 5.72 | 5.65 | 4.75 | 89.3±0.42 | 60.63±1.9 |
| **8** | **1** | 5.7 | 5.3 | 0 | 100 | |
| | **2** | 5.7 | 5.6 | 0 | 100 | |
| | **3** | 5.7 | 5.6 | 0 | 100 | |
| Average: | | 5.72 | 5.52 | 0 | 100.00±0.00 | |
| **24** | **1** | 5.04 | 4.8 | 0 | 100 | |
| | **2** | 501 | 4.8 | 0 | 100 | |
| | **3** | 5.07 | 4.9 | 0 | 100 | |
| Average: | | 5.09 | 4..88 | 0 | 100.00±0.00 | |

Camera images and their CFU/ml estimates of the *E. coli* and *E. faecalis* bacteria colonies are depicted in **Figure S5**.

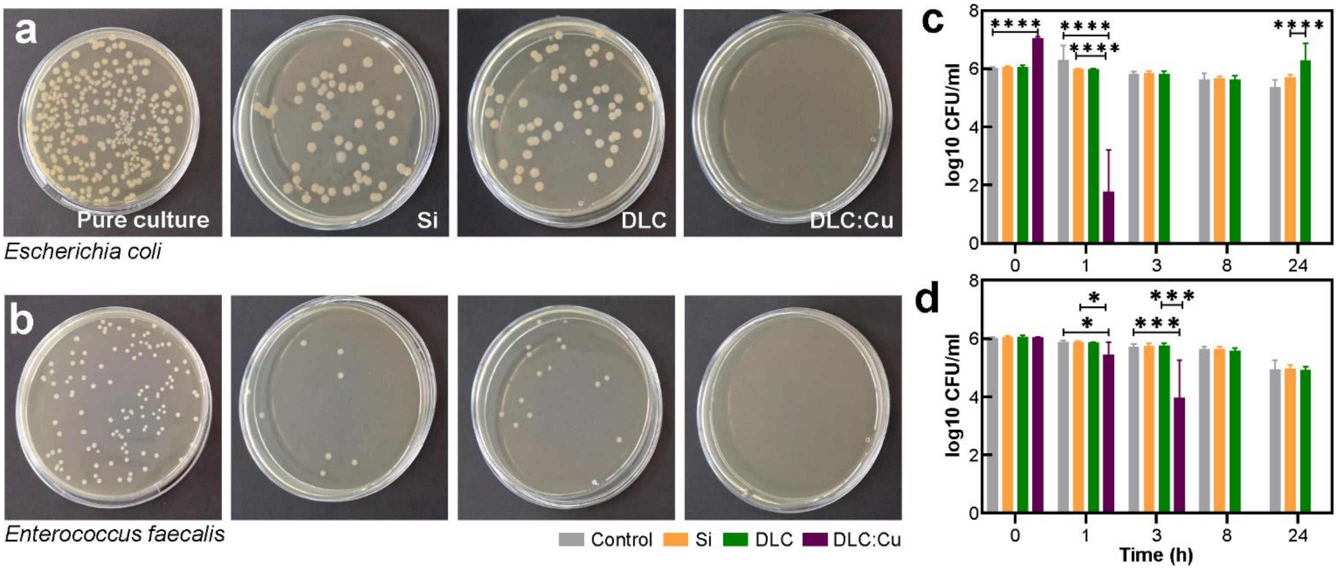


**Figure S5** Camera images of bacteria colonies grown on agar plates for Si, DLC, and DLC:Cu obtained after 24 hour immersion in (a) *E. coli* and (b) *E. faecalis* while the control was pure culture without any treatment. The $\log_{10}$ CFU/ml of both investigated bacteria is shown in the figures (c) and (d), respectively.

### References


(1) Takata, S.; Wada, H.; Tamura, M.; Koide, T.; Higaki, M.; Mikura, S.; Yasutake, T.; Hirao, S.; Nakamura, M.; Honda, K.; et al. Kinetics of c-reactive protein (CRP) and serum amyloid A protein (SAA) in patients with community-acquired pneumonia (CAP), as presented with biologic half-life times. *Biomarkers* **2011**, *16* (6), 530-535, Article. DOI: 10.3109/1354750x.2011.607189.

(2) Biesinger, M. C.; Lau, L. W. M.; Gerson, A. R.; Smart, R. S. C. Resolving surface chemical states in XPS analysis of first row transition metals, oxides and hydroxides: Sc, Ti, V, Cu and Zn. *Applied Surface Science* **2010**, *257* (3), 887-898, Article. DOI: 10.1016/j.apsusc.2010.07.086.

(3) NIST X-ray Photoelectron Spectroscopy Database, NIST Standard Reference Database Number 20. National Institute of Standards and Technology: Gaithersburg MD, 2000.

(4) Biesinger, M. C. Advanced analysis of copper X-ray photoelectron spectra. *Surface and Interface Analysis* **2017**, *49* (13), 1325-1334, Article. DOI: 10.1002/sia.6239.
